# Microglia associations with brain pericytes and the vasculature are reduced in Alzheimer’s disease

**DOI:** 10.1101/2022.08.08.503250

**Authors:** Gary P. Morris, Catherine G. Foster, Jo-Maree Courtney, Jessica M. Collins, Jake M. Cashion, Lachlan S. Brown, David W. Howells, Gabriele C. DeLuca, Alison J. Canty, Anna E. King, Jenna M. Ziebell, Brad A. Sutherland

## Abstract

Cerebral blood flow is important for the maintenance of brain function and its dysregulation has been implicated in Alzheimer’s disease (AD). Subpopulations of microglia have well-characterised associations with the vasculature in the central nervous system but the precise relationship between microglia and cells which exist on the vasculature is not yet clear. In this study we explored the relationship between microglia and pericytes, a vessel-resident cell type that has a major role in the regulation of cerebral blood flow and maintenance of the blood brain barrier. Using fixed tissue sections and *in vivo* live imaging, we discovered a subset of microglia that closely associated with pericytes, termed <u>PE</u>ricyte-associated <u>M</u>icroglia (PEM). PEM are present throughout all regions of the brain and spinal cord in NG2DsRed x CX_3_CR1^+/GFP^ mice, and in the human frontal cortex. They reside adjacent to pericytes at all levels of the capillary tree and can maintain their position for at least 28 days. PEM associate with pericytes lacking astroglial endfeet coverage but are segregated from pericytes by capillary basement membranes and capillary vessel width is similarly increased beneath pericytes with or without an associated PEM. Deletion of the microglia fractalkine receptor (CX_3_CR1) did not disrupt the association between pericytes and PEM, suggesting the association is not reliant on fractalkine signalling. Finally, we found that the proportion of microglia that are capillary-associated and PEM declines in the superior frontal gyrus (SFG) in AD, which is exacerbated by the *APOE* ε3/ε4 genotype. In summary, we identify and characterise a subpopulation of microglia that specifically associate with pericytes and find this population is reduced in the SFG in AD. This reduction may be a novel mechanism contributing to vascular dysfunction in diseases such as AD.

## Introduction

Microglia are highly ramified, dynamic cells that contribute to homeostatic functions including structural plasticity, synaptic plasticity, neurite formation, myelination and vasculogenesis^1-3^. Microglia are also central players in the innate immune system of the central nervous system (CNS), rapidly responding to challenges to the CNS and playing important roles in the initiation of inflammation, removal of debris and promotion of various CNS recovery mechanisms^3^. A subpopulation of microglia are known to have a close relationship with the vasculature during CNS development and in adulthood^4-8^, with these microglia having been variously termed “vascular satellites^7,9,10^”, “perivascular microglia^7,9,11^”, “juxtavascular microglia^9,12,13^”, “vessel-associated microglia^14^”, and more recently “capillary-associated microglia (CAM)^15^”. Given a subset of microglia reside adjacent to vessels, microglia may play a role in blood vessel function and maintenance. In support of this, recent studies have illustrated that ablation of microglia triggers capillary dilation^15^ and impairs evoked blood flow changes following stimulation^15,16^. The mechanisms governing their contribution to these functions are not yet clear, but it is possible they influence blood flow indirectly through pericytes^15,17^. Pericytes are cells embedded on the walls of capillaries throughout the body^18,19^ and are well characterised as modulators of cerebral blood flow^20^. However, little is known about the relationship between microglia and pericytes. Early reports confused the two cell types due to their close spatial relationship^21^ and a collection of evidence suggests pericytes may differentiate into microglia^7,22,23^. Outside of these findings, it is not known how often microglia and pericytes are spatially associated, or whether any spatial association between the two cell types may serve a functional purpose.

As many as 80% of individuals diagnosed with Alzheimer’s disease (AD) have some level of vascular pathology in the brain including microinfarcts and cerebral amyloid angiopathy^24^. Both impaired cerebral blood flow and blood-brain barrier breakdown are also reported to occur early in the pathogenesis of AD^25^. Although the aetiology of vascular alterations in AD is unknown, increasing evidence has implicated pericytes in blood flow dysfunction, neurodegeneration and cognitive decline in AD^26,27^. In the human brain, several studies have suggested pericytes are susceptible to death in brain regions affected by AD^28-30^, which would hypothetically have detrimental consequences for blood flow regulation. Similarly, a growing body of literature has implicated microglia dysfunction as a key feature of AD pathogenesis. Several sporadic AD risk genes are highly expressed in microglia^31,32^, maladaptive microglia over-strip synapses in mouse models of AD^33^ and sub-populations of microglia adopt specific phenotypes in AD^34^. Given the role pericytes and microglia may play in the pathogenesis of AD, any breakdown in a functional association between these two cell types may also influence AD pathogenesis.

In this study, we first investigated the spatial relationship between microglia and pericytes in the adult mouse brain and spinal cord. In doing so, we discovered a subset of microglia that dynamically interact with pericytes. We have named these cells pericyte-associated microglia (**PEM**). Using *in vivo* two-photon laser scanning microscopy (2PLSM), we found that PEM can maintain their position adjacent to pericytes for at least 28 days in the somatosensory cortex and are present at all levels of the capillary vascular tree. In post-mortem human AD tissue, we found a lower proportion of microglia that were PEM in the frontal cortex compared to controls, despite an increase in the number of pericytes, an effect that was exacerbated in *APOE4* carriers. These results point towards a close relationship between microglia and pericytes that, if disrupted, could contribute to the vascular pathogenesis of neurodegenerative diseases, including AD.

## Results

### A subset of microglia are adjacent to capillary pericytes in the healthy mouse brain and spinal cord

Given that there has not yet been a formal investigation of the spatial relationship between microglia and pericytes, we first determined if microglia and pericytes reside adjacent to one another and how often such associations occur in the healthy brain. To do this, we generated NG2DsRed x CX_3_CR1^+/GFP^ mice to enable robust visualisation of microglia (CX_3_CR1^+/GFP^) and pericytes (NG2DsRed) in fixed brain tissue (Fig. 1A-B).

**Figure 1.**
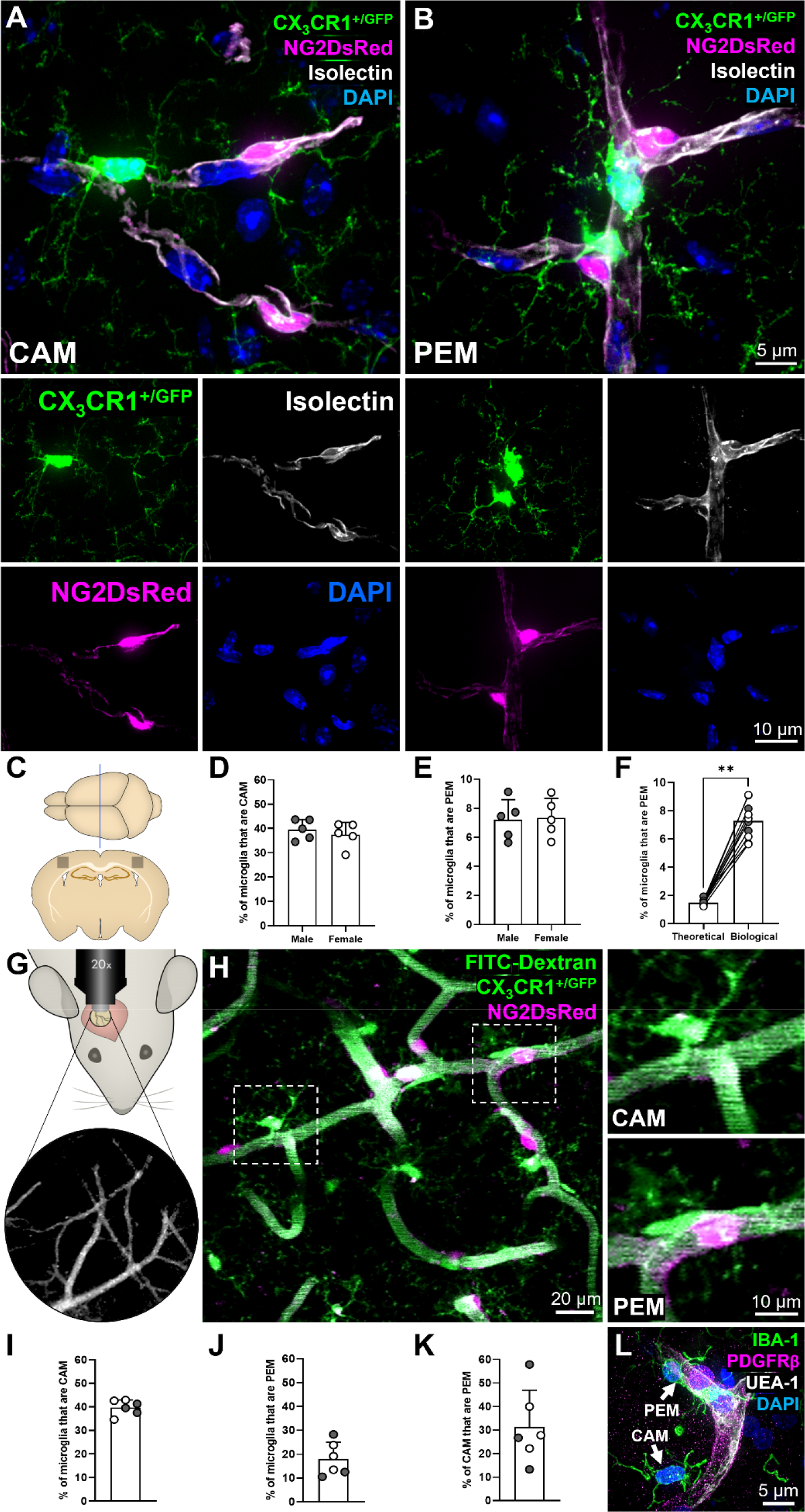
A subset of microglia are directly adjacent to pericytes. **(A)** Representative example of a capillary-associated microglia (CAM), in the somatosensory cortex, extending fine processes to contact other DAPI-positive nuclei. **(B)** Representative example of two pericyte-associated microglia (PEM), in the hippocampus, with the bottom PEM morphologically curved around a pericyte. (A-B) For all images: NG2DsRed-positive pericytes (magenta), CX_3_CR1^+/GFP^-positive microglia (green), isolectin-labelled vessels (white) and DAPI-labelled nuclei (blue). Each fluorescent channel alone is located below the main image. All images were derived from 12-week-old NG2DsRed x CX_3_CR1^+/GFP^ mice using confocal microscopy. **(C)** Schematic of region analysed within the somatosensory cortex of NG2DsRed x CX_3_CR1^+/GFP^ mice (Bregma -1.5 mm^67^). **(D-E)** Quantification of (D) CAM and (E) PEM in the somatosensory cortex of male (*n* = 5) and female (*n* = 5) mice. Data compared with an unpaired parametric t-test. **(F)** The predicted percentage of PEM when running simulations with microglia and pericyte densities derived from ten different biological replicates, compared to the actual PEM percentage from those same ten somatosensory biological replicates (biological data derived from Fig. 1E). Data compared with a Wilcoxon test. Lines represent paired data. **(G)** Schematic of cranial window location in NG2DsRed x CX_3_CR1^+/GFP^ mice with blood vessels used as landmarks. **(H)** Representative 30 μm thick projection image of NG2DsRed-positive pericytes (magenta), CX_3_CR1^+/GFP^-positive microglia (green) and FITC-dextran-positive vessel lumen (green) in layers II/III of the somatosensory cortex of adult NG2DsRed x CX_3_CR1^+/GFP^ mice imaged using 2PLSM. Dashed boxes highlighting a CAM and PEM are magnified in panels to the right. **(I-K)** Quantification of the percentage of microglia that are (I) CAM, (J) PEM and (K) the percentage of CAM that are PEM, in layers II/III of the somatosensory cortex of male (*n* = 3) and female (*n* = 3) mice. **(L)** Representative image of PDGFRβ-positive pericyte (magenta), IBA1-positive microglia (green), UEA-1-labelled vessels (white) and DAPI-labelled nuclei (blue) from the SFG of a human control brain (93 y.o. female, *APOE* ε3/ε3). A CAM and PEM are highlighted by white arrows. For all graphs, grey circles represent males and white circles represent females. Data presented as mean ± SD. ** *p* < 0.01.

Confocal microscopy of brain tissue sections highlighted two sub-populations of microglia. One subpopulation closely associates with capillaries, which we refer to as capillary-associated microglia (CAM, Fig. 1A, Supplementary Movie 1) consistent with a previous report^15^. The second is a newly identified subpopulation of microglia that reside directly adjacent to pericytes on capillaries (Fig. 1B, Supplementary Fig. 1, Supplementary Movies 2-3). We have termed these <u>pe</u>ricyte-associated <u>m</u>icroglia (PEM). Both CAM and PEM were observed extending processes toward other nuclei, suggesting they may make multiple contacts with blood vessels, pericytes and other cell soma in the brain parenchyma simultaneously (Fig. 1A-B, Supplementary Fig. 1, Supplementary Movies 1-3). We also frequently observed PEM soma exhibiting a curved morphology around pericyte cell bodies (Fig. 1B, L, Supplementary Fig. 1, Supplementary Movies 2-3, Fig. 3B) and the processes of both PEM and CAM reaching out to enfold or contact pericyte cell bodies (Fig. 1B, Fig. 3A, C).

Quantification revealed a large proportion of microglia were CAM (male = 39.6 ± 4.2%, female = 37.5 ± 5.1%, Fig. 1C-D), which is consistent with previous reports^15^. Next, we assessed the proportion of microglia that were PEM, finding in tissue slices ∼1 in 15 microglia were located adjacent to pericytes (male = 7.2 ± 1.4%, female = 7.3 ± 1.3%, Fig. 1C, E). There were no significant differences in the proportion of microglia that were CAM or PEM between male and female mice (Fig. 1D-E). To determine whether the proportion of microglia that were PEM was higher than would be expected by random chance, we ran simulations to mathematically model chance associations between pericytes and microglia. We found the percentage of PEM to be ∼5 fold more than expected by chance (7.3 ± 1.3% vs. 1.5 ± 0.19%, Fig. 1F, Supplementary Fig. 2).

To determine if PEM are a ubiquitous feature of the CNS, we expanded our analysis of microglia-pericyte associations to also include the caudate putamen, hippocampus, thalamus, hypothalamus and the spinal cord (Supplementary Fig. 3). The proportion of microglia that were identified as PEM was consistent across all regions assessed (Supplementary Fig. 3B-D), though there were some differences in the number of pericytes and microglia and the proportion of pericytes with a PEM (Supplementary Fig. 3E-G). Furthermore, we identified the presence of both CAM and PEM in the thoracic region of the spinal cord of NG2DsRed x CX_3_CR1^+/GFP^ mice (Supplementary Fig. 3H-L).

We next replicated and extended this analysis by quantifying the proportion of microglia that were CAM and PEM in the upper layers of the somatosensory cortex (layers II/III), using images derived from *in vivo* 2PLSM taken through cranial windows (Fig. 1G). This approach allowed us to assess the spatial relationship between pericytes and microglia in three dimensions through large z-stacks (∼130 μm in depth, Fig. 1H, Supplementary Movie 4). Similar to our analysis in fixed tissue, we found that ∼2 in every 5 microglia were CAM (39.7 ± 3.2%, Fig. 1I), but that ∼1 in every 6 microglia were PEM (18.0 ± 7.0%, Fig. 1J), a higher proportion than in our fixed tissue analysis. Regions were manually selected for *in vivo* imaging if they were specifically enriched in PEM, likely partially explaining this difference. Of the microglia that were CAM, ∼1/3^rd^ of them also met our definition of PEM (31.3 ± 15.7%, Fig. 1K). We also identified both PEM and CAM in the SFG of post-mortem human tissue (Fig. 1L, Supplementary Movie 5), illustrating they are conserved across species.

Given that pericytes reside throughout the entire capillary bed, we used the *in vivo* 2PLSM images to determine if CAM and PEM preferentially reside at specific levels of the vascular tree (Fig. 2, Supplementary Fig. 4). CAM, PEM and pericytes were found at every level of the vascular tree (Fig. 2B), at both vessel junctions (Fig. 2C) and straight vessel segments (Supplementary Fig. 4C), with no clear positional preference. Collectively, these findings illustrate that a subset of microglia are directly adjacent to pericytes in the mouse brain and spinal cord, as well as the human brain and that they are evenly distributed throughout the vascular tree.

**Figure 2.**
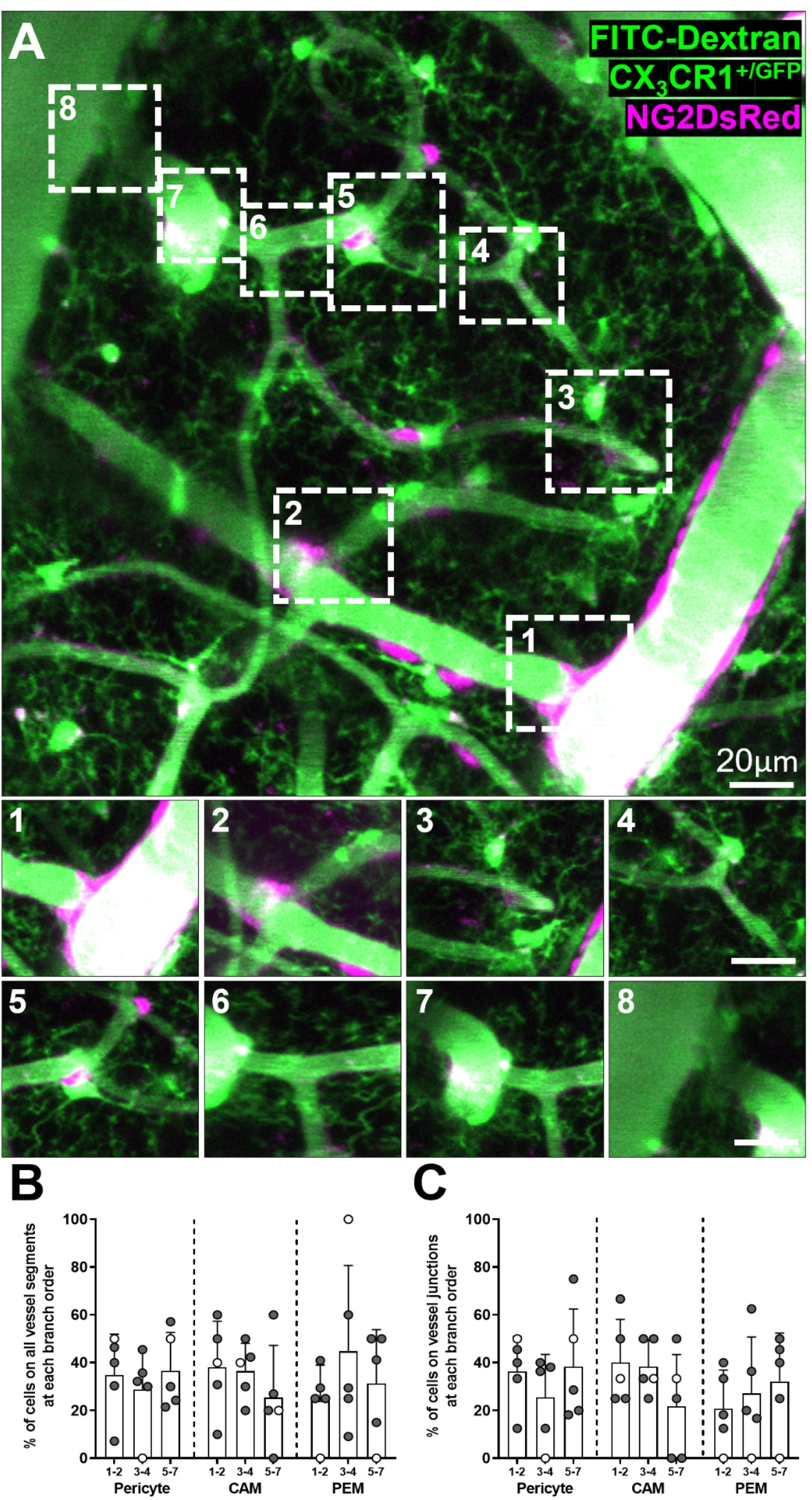
CAM, PEM and pericytes are present at all branch orders of capillaries. **(A)** Representative image stack (20-75μm depth, average intensity projection) of FITC dextran filled vessels in the somatosensory cortex of an NG2DsRed x CX_3_CR1^+/GFP^ mouse imaged *in vivo* using 2PLSM. Pericytes are labelled magenta, and microglia and vessels are green. Insets match boxes labelled 1-8 in main panel. (1) Penetrating arteriole (0th order) branching off to form a capillary (first order). (2-7) Higher order capillaries branching. (8) Seventh order capillary converging on the ascending venule. **(B-C)** Percentage of total CAM, PEM and pericytes that are located (B) at different branch orders of the vascular tree, and (C) at vessel junctions at different branch orders of the vascular tree (*n* = 5, four male and one female). For all graphs, grey circles represent males and white circles represent females. Data presented as mean ± SD.

### Microglia associate with pericytes lacking astrocyte endfeet coverage

It has previously been demonstrated that microglia associate with vessels lacking astrocyte endfeet coverage^13,15^. We confirmed this by showing CAM contacting blood vessels lacking AQP4 labelling in the somatosensory cortex of 12-week-old NG2DsRed x CX_3_CR1^+/GFP^ mice (Fig. 3A). Similarly, we found that where PEM are adjacent to pericytes, pericytes can lack complete astrocyte endfeet coverage (Fig. 3B-C, Supplementary Movies 6-7). 3D reconstructions (Fig. 3D, Supplementary Movies 6-8) and depth shading (Fig. 3E) illustrate that microglial cell bodies and processes associate with pericytes lacking AQP4 labelling. These reconstructions also reveal gaps between the associating microglia and pericytes, despite the lack of AQP4 labelling between them (Fig. 3D-E and Supplementary Movies 7-8). Staining of the basement membrane with isolectin revealed these gaps can be filled by basement membrane (Supplementary Fig. 5, Supplementary Movies 9-10). Together, these findings illustrate that PEM likely reside outside of the basement membrane, but inside the AQP4-positive astrocyte endfeet layer.

**Figure 3.**
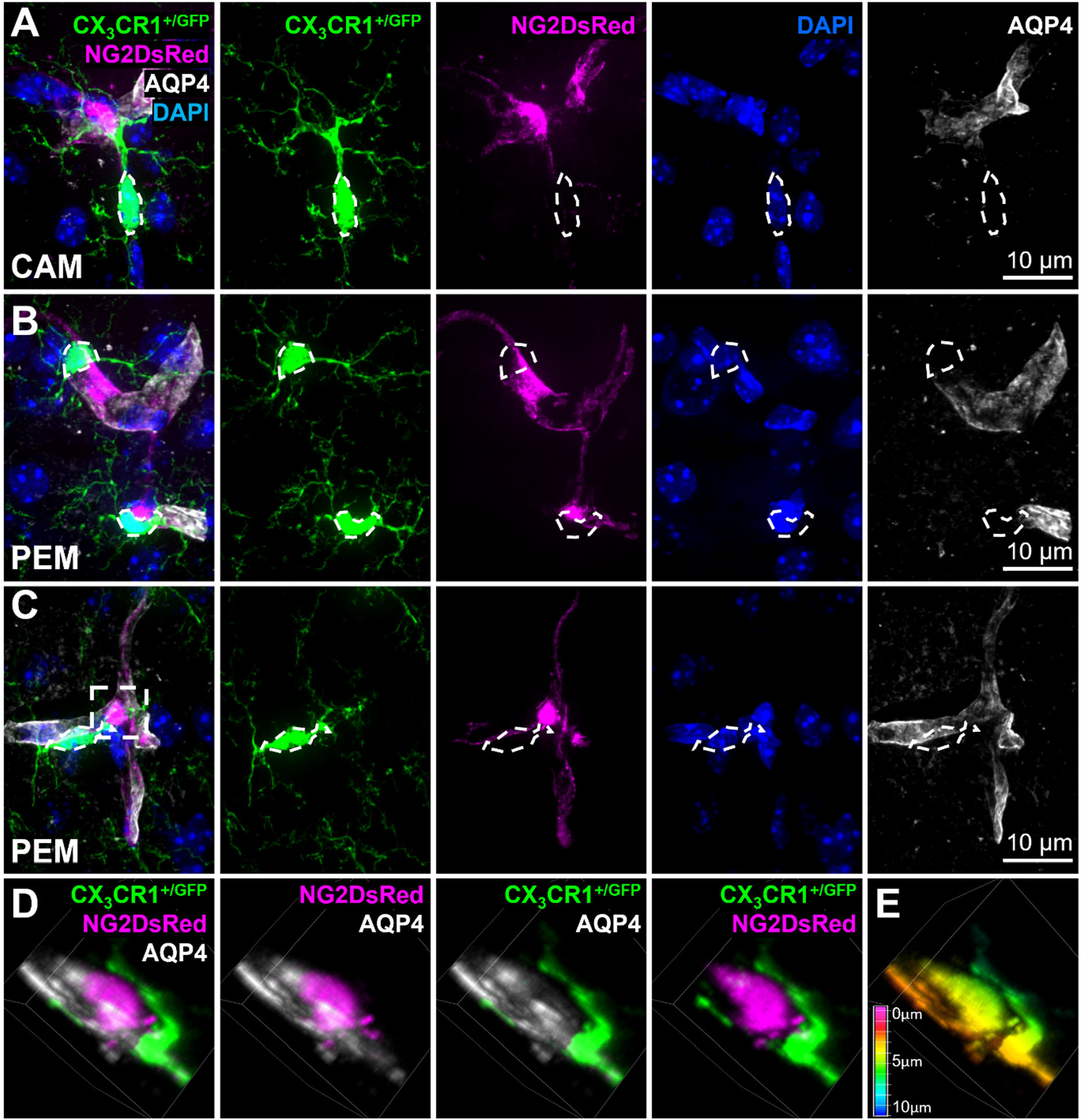
Pericyte-associated microglia interact with pericytes lacking AQP4 coverage. **(A)** Representative example of a CAM from the somatosensory cortex residing on a vessel lacking AQP4 coverage. Maximum projection image from a 24 μm z-stack, imaged with 1 μm increments. **(B)** Representative example of two PEM, one of which is morphologically curved around a pericyte in the somatosensory cortex on a blood vessel lacking AQP4 coverage. **(C)** Representative example of AQP4 coverage lacking at the site in which the process of a microglia is reaching around a pericyte in the somatosensory cortex. (A-C) For all images: NG2DsRed-positive pericytes (magenta), CX_3_CR1^+/GFP^-positive microglia (green), AQP4-labelled astrocyte endfeet (white) and DAPI-labelled nuclei (blue) are shown. Images showing each fluorescent channel alone are to the right of the main image. **(D)** Magnification of dashed box in (C), rotated to illustrate the association between the pericyte soma and a microglial process at an area lacking AQP4 labelling. Each panel is rotated identically. **(E)** Depth shading of image in (D) illustrating the microglia process is in the same plane as the pericyte but AQP4 labelling is below the plane of the pericyte and microglia process. All images were derived from 12-week-old NG2DsRed x CX_3_CR1^+/GFP^ mice using confocal microscopy.

### Microglial associations with pericytes can be gained and lost

To determine if PEM maintain their position adjacent to pericytes over time, we tracked 32 PEM over 28 days, through cranial windows in NG2DsRed x CX_3_CR1^+/GFP^ mice (Supplementary Fig. 6). Throughout the imaging period, we identified PEM that fell within three groups: PEM that remained adjacent to their partner pericytes over the entire 28-day period (Fig. 4A); PEM that were no longer adjacent to a pericyte within the 28 days (Fig. 4B); and pericytes that did not originally have a PEM, but which gained one during the 28-day period (Fig. 4C). Upon quantification, we identified that the majority of PEM from day 0 were still positioned directly adjacent to a pericyte after four days (81%) and seven days (78%), but by day 28, only 44% of PEM remained adjacent to a pericyte (Fig. 4D). Despite this loss of PEM, we did not observe a change in the total proportion of microglia that were defined as a PEM throughout the 28 days, indicating some pericytes that did not originally have a PEM had gained a PEM during the imaging period (Fig. 4E). Collectively, these data suggest microglial associations with pericytes are dynamic, are consistent in proportion over time, and only a minority of PEM have associations with pericytes that last 28 days.

**Figure 4.**
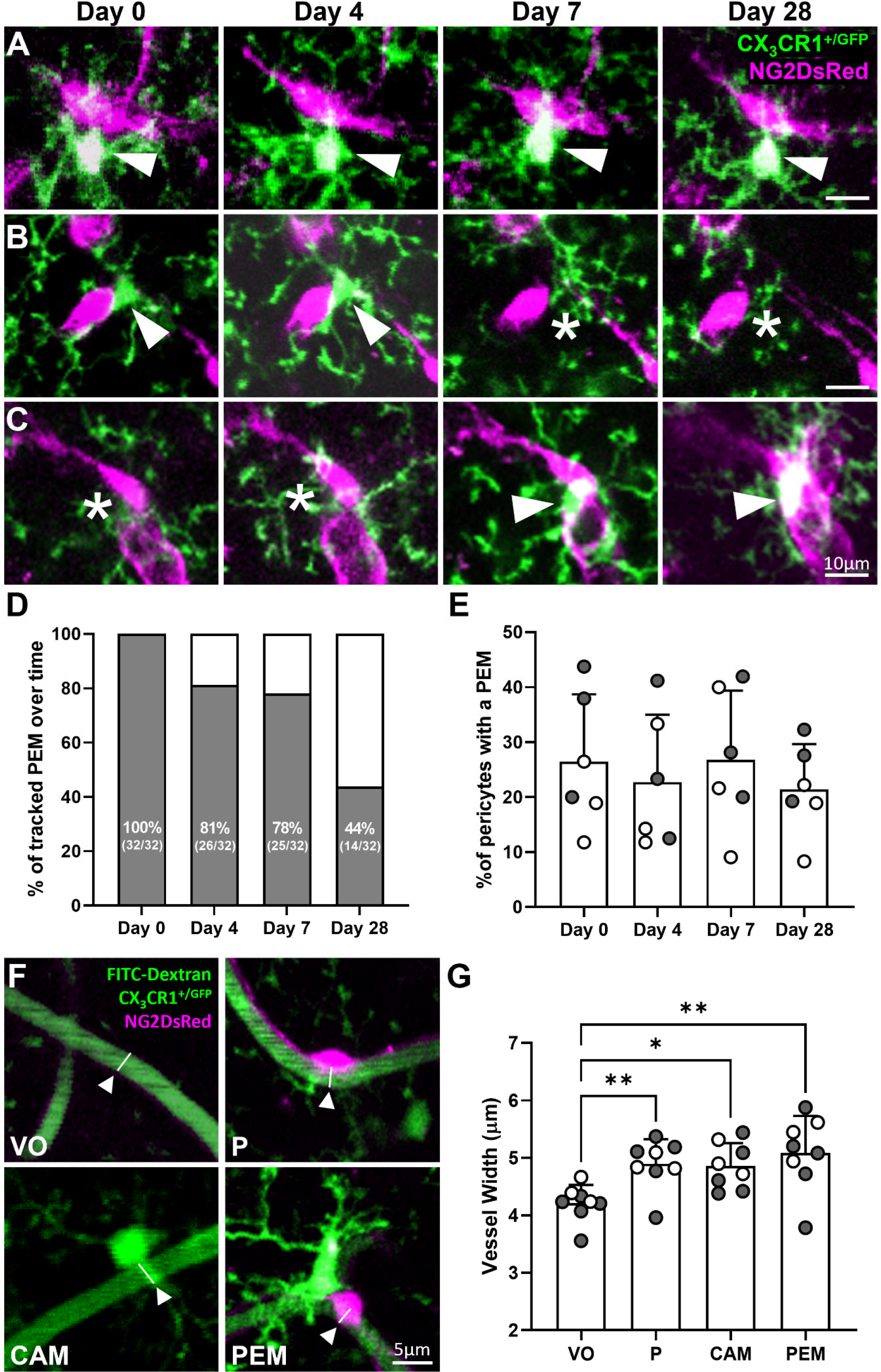
Pericytes can gain or lose PEM over 28 days and vessel width is increased beneath pericyte and microglia soma. **(A)** Representative *in vivo* 2PLSM images of a PEM that maintains position for 28 days in the somatosensory cortex of an NG2DsRed x CX_3_CR1^+/GFP^ mouse. **(B)** Representative images of a PEM in position for 4 days that is no longer in place on day 7 or 28 of imaging. **(C)** Representative images of a pericyte with no adjacent PEM on day 0 or 4 but gains a PEM on day 7. (A-C) For all images pericytes are magenta, microglia are green, and images are an average intensity projection. White arrowheads highlight a PEM and white asterisks indicate the absence of a PEM. **(D-E)** Percentage of (D) 32 PEM, identified on day 0 of imaging, that maintain position adjacent to a pericyte on days 4, 7 and 28 of imaging and (E) total pericytes with a PEM per mouse on days 0, 4, 7 and 28 (*n* = 6, three male and three female). Data in (E) analysed using a repeated measures one-way ANOVA. (**F)** Representative images indicating locations of vessel width measurement in *in vivo* 2PLSM images derived from NG2DsRed x CX_3_CR1^+/GFP^ mice, following administration of the vessel lumen marker FITC-dextran. White arrowheads indicate the midpoint below the cell soma. **(G)** Quantification of vessel width (μm) of vessel only (VO), and vessels with a pericyte (P), CAM or a pericyte with a PEM (150, 124, 90 and 124 vessels were measured, respectively, across *n* = 8 mice, five male and three female). Data presented as average per animal with a minimum of four vessel measurements at each location. Comparisons were made with a Friedman’s test. For all graphs, grey circles represent males and white circles represent females. Data presented as mean ± SD. \**p* < 0.05, \*\**p* < 0.01.

### Capillary width is increased beneath the soma of microglia and pericytes with or without a PEM

To investigate if CAM or PEM may influence blood flow by altering capillary diameter in the healthy brain, we measured capillary width beneath pericytes, CAM and pericytes with a PEM, compared to vessel segments without any observable associated pericytes, CAM or PEM (Fig. 4F). We found that the average vessel width for vessels with no observable associated cells was 4.21 ± 0.32 μm (Fig. 4G). Compared to vessels with no observable cells (vessel only, VO), the vessel width was significantly increased by 16% beneath pericytes (4.89 ± 0.43 μm, *p* = 0.0060) and by 15% beneath CAM (4.86 ± 0.40 μm, *p* = 0.0402). Beneath pericytes with an adjacent PEM, the width of vessels was 21% wider (5.08 ± 0.64 μm, *p* = 0.0060) than vessels with no observable cells (Fig. 4G). These findings suggest the presence of both pericytes and microglia on capillaries is associated with increased capillary width under basal conditions, but that the presence of a PEM near a pericyte does not alter this.

### CX_3_CR1 knock-out does not alter the proportion of microglia that are CAM or PEM

The signals that recruit microglia to pericytes are unknown. Microglia migrate in response to the fractalkine ligand (CX_3_CL1), which can be released from neurons and binds to the fractalkine receptor (CX_3_CR1) on microglia. Recently, CX_3_CL1 was shown to cause capillary constriction at regions where microglia contact vessels, in both the brain and retinal explants, suggesting the CX_3_CL1/ CX_3_CR1 axis may play a role in microglia modulation of vessel tone^35^. Pericytes also have the capacity to release CX_3_CL1^36^, so it is possible they may be a source of CX_3_CL1 to recruit microglia to the vasculature. To determine if microglia are recruited to vessels and pericytes through the CX_3_CL1/CX_3_CR1 axis, we took advantage of the CX_3_CR1^GFP^ knock-in/knock-out line, which replaces the first 390 bp of exon 2 of CX_3_CR1 with GFP. We derived both the single KO NG2DsRed x CX_3_CR1^+/GFP^ line and the complete KO NG2DsRed x homozygous CX_3_CR1^GFP/GFP^. Confirmation of homozygosity was through genotyping and confirmed by the observation of increased GFP fluorescence intensity in CX_3_CR1^GFP/GFP^ microglia, compared to CX_3_CR1^+/GFP^ microglia (Supplementary Fig. 7A-D). There was no significant difference in the number of microglia or pericytes in the somatosensory cortex of 12-week-old NG2DsRed x CX_3_CR1^+/GFP^ mice compared to NG2DsRed x CX_3_CR1^GFP/GFP^ mice (Supplementary Fig. 7E-F). Both CAM and PEM were present in both heterozygous and homozygous KO mice (Fig. 5A-B), and upon quantification there was no significant difference in the proportion of microglia that were CAM or PEM in heterozygous vs. homozygous KOs (CAM: heterozygous 42.3 ± 3.3% vs. homozygous 43.6 ± 2.8%, Fig. 5C; PEM: heterozygous 6.9 ± 1.1 vs. homozygous 7.4 ± 0.8 PEM respectively, Fig. 5D).

**Figure 5.**
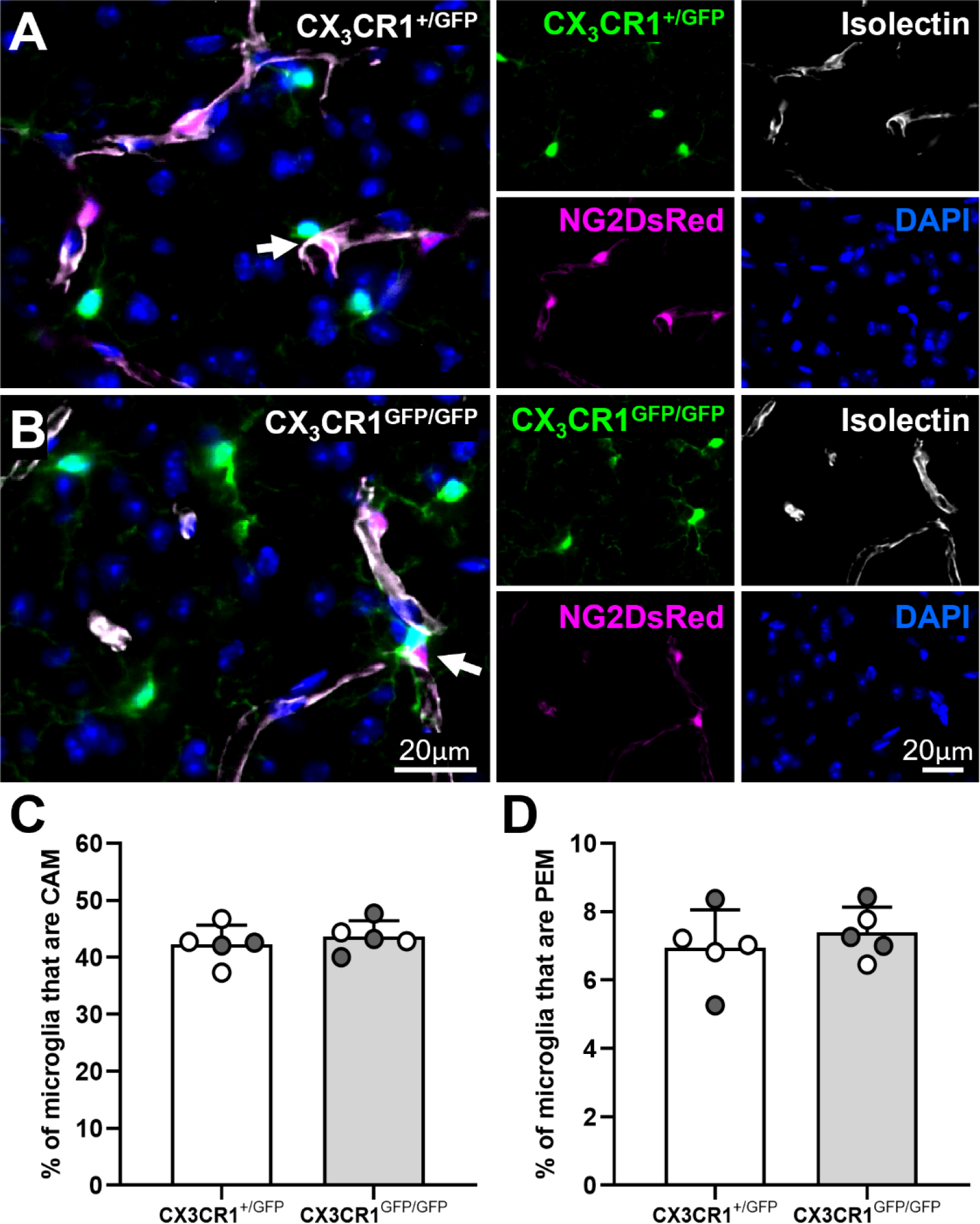
CX_3_CR1 knock-out does not alter the proportion of microglia that are PEM or CAM in 12-week-old mice. **(A-B)** Representative images of (A) NG2DsRed x CX_3_CR1^+/GFP^ and (B) NG2DsRed x CX_3_CR1^GFP/GFP^ mice showing that PEM are present in both mouse lines (white arrows). For each image: NG2DsRed-positive pericytes (magenta), CX_3_CR1^+/GFP^-positive microglia (green), AQP4-labelled astrocyte endfeet (white) and DAPI-labelled nuclei (blue) are shown. Images showing each fluorescent channel alone are to the right of the main image. **(C-D)** Quantification of (C) CAM and (D) PEM in the somatosensory cortex (Bregma -1.5; *n* = 5 per group, two males and three females for NG2DsRed x CX_3_CR1^+/GFP^, three males and two females for NG2DsRed x CX_3_CR1^GFP/GFP^). Data compared with an unpaired parametric t-test. For all graphs, grey circles represent males and white circles represent females. Data presented as mean ± SD.

### CAM and PEM are reduced in the Alzheimer’s disease superior frontal gyrus

To assess whether there is a change in the prevalence of microglia and pericyte associations in AD, we assessed CAM, PEM, microglia and pericytes in post-mortem tissue from the SFG of a human AD cohort. Both microglia and pericytes could be visualised in the human brain and both CAM and PEM were identifiable in control and AD tissue (Figs. 1L, 6A-C, Supplementary Movies 5, 11). Quantification of microglia found no significant difference in the number of microglia in controls vs. AD (control 105.2 ± 21.2 vs. AD 119.8 ± 31.4 per mm^2^, Fig. 6D). However, we observed a significant increase in the number of pericytes in the SFG of AD cases, compared to controls (control 71.9 ± 16.0 vs. AD 89.9 ± 17.9 per mm^2^, *p* = 0.0336, Fig. 6E). Quantification of the proportion of microglia that were CAM and PEM showed they were present in similar proportions to mouse brain tissue sections (∼30-40% of microglia were CAM and ∼4-5% of microglia were PEM, Figs. 6F-G). Despite the increased density of pericytes throughout the SFG in the AD brain, we observed a significant decrease in the proportion of microglia that were CAM and PEM in AD compared to controls (CAM: control 37.7 ± 7.6% vs. AD 28.2 ± 2.7%, *p* = 0.0017, Fig. 6F; PEM: control 4.9 ± 1.9% vs. AD 3.4 ± 1.1%, *p* = 0.039, Fig. 6G). Finally, we assessed microvessel density, observing no difference in the overall length of microvessels in AD brain sections compared to control brain sections (control 7559 ± 2197 vs. AD 8528 ± 2199 μm/mm^2^, *p* = 0.30, Supplementary Fig. 8A).

**Figure 6.**
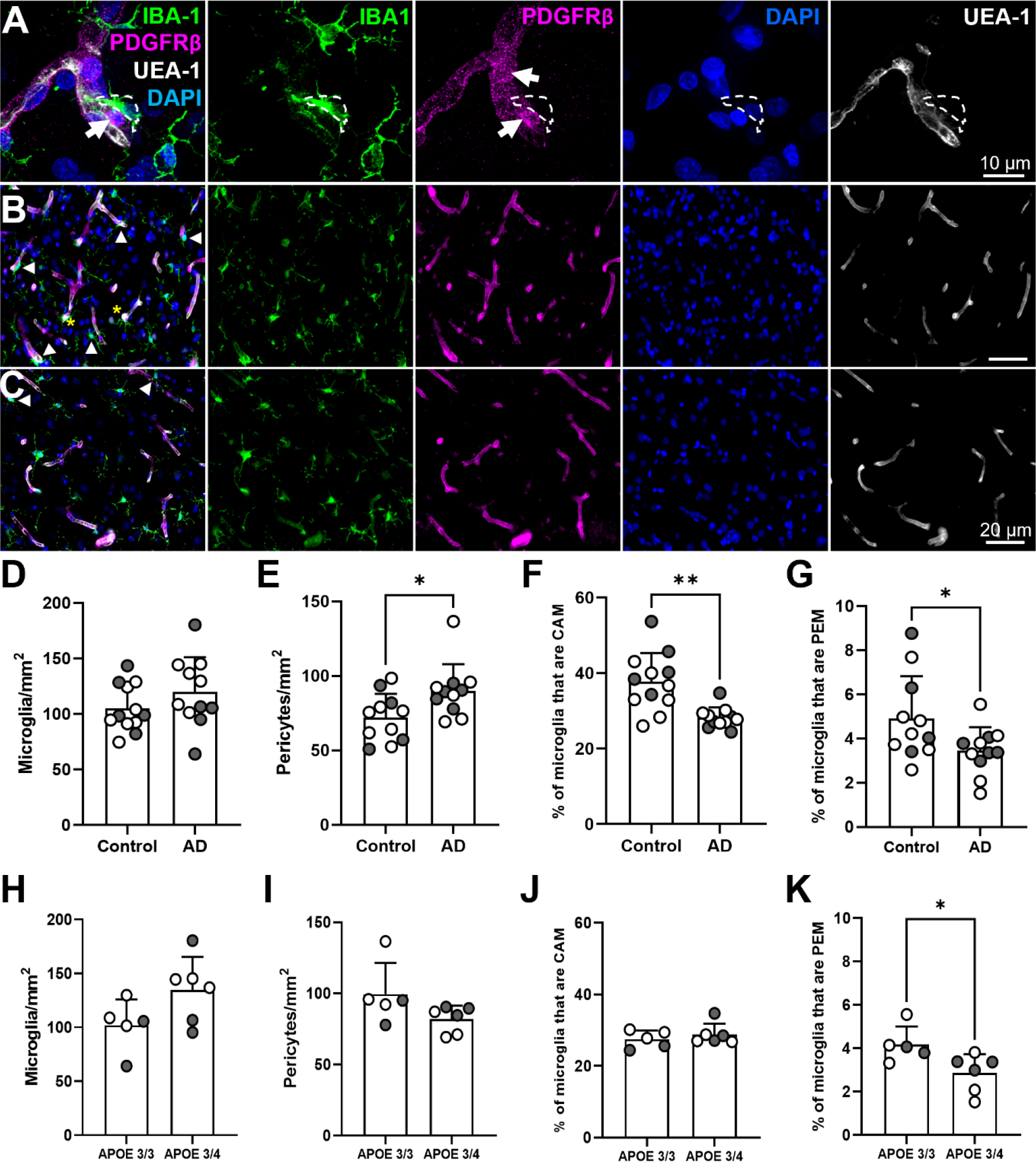
There are more pericytes in the superior frontal gryus in Alzheimer’s disease, but the proportion of microglia that are PEM and CAM is reduced. **(A)** Representative image of PDGFRβ-positive pericytes (magenta), IBA1-positive microglia (green), UEA-1-labelled vessels (white) and DAPI-labelled nuclei (blue) from SFG of a human control brain (93 y.o. female, *APOE* ε3/ε3). IBA1, PDGFRβ, DAPI and UEA-1 channels are shown alone to the right of main image. White arrows identify pericyte soma. White dashed lines outline the position of a PEM. **(B-C)** Representative images of IBA1, PDGFRβ, DAPI and UEA-1 combined, and split into individual channels, in the SFG of (B) a control brain and (C) an AD brain. White arrowheads denote CAM and yellow asterisks denote PEM. **(D-G)** Quantification of (D) microglia, (E) pericytes, (F) CAM and (G) PEM in the SFG of control (*n* = 11-12) and AD (*n* = 11) post-mortem brains. (D, G) Analysed with an unpaired parametric t-test. (E-F) Analysed with an unpaired nonparametric Mann-Whitney test. **(H-K)** Quantification of (H) microglia, (I) pericytes, (J) CAM and (K) PEM in *APOE* ε3/ε3 (*n* = 5) and *APOE* ε3/ε4 (*n* = 6) in AD. (H-I, K) Analysed with an unpaired parametric t-test. (J) Analysed with an unpaired nonparametric Mann-Whitney test. For all graphs, grey circles represent males and white circles represent females. Data are presented as mean ± SD. \**p* < 0.05, \*\**p* < 0.01.

We next segregated the AD cohort based on *APOE* genotype status (*APOE* ε3/ε3 *n* = 5, *APOE* ε3/ε4 *n* = 6). *APOE* status had no effect on the number of microglia, pericytes and proportion of microglia that were CAM (Fig. 6H-J), but there was a significant decrease in the proportion of microglia that were PEM in AD cases with an *APOE* ε3/ε4 genotype, compared to those with an *APOE* ε3/ε3 genotype (ε3/ε3 4.2 ± 0.8% vs. ε3/ε4 2.9 ± 0.9%, *p* = 0.0322, Fig. 6K). Collectively, these data suggest that the SFG in AD is characterised by an increase in pericyte density, a reduction in the association of microglia with capillaries and pericytes, and that the reduction in PEM is exacerbated by the presence of *APOE* ε3/ε4 genotype.

## Discussion

We identify and describe a unique subset of microglia that dynamically interact with capillary pericytes in the mouse brain and spinal cord, and the human brain. For simplicity, we have named these cells pericyte-associated microglia (PEM). Collectively, our data, combined with reports of similar interactions in the retina^35^, suggest PEM are an ubiquitous feature of the CNS present at all levels of the capillary tree.

PEM likely overlap with other subtypes of microglia. In particular, because we focused our analysis specifically on capillaries <10 μm in diameter, the PEM we analysed most closely overlap with CAM^15^. Our data demonstrate CAM are unlikely to be preferentially adjacent to pericytes, as the proportion of microglia that are CAM outnumber the proportion of microglia that are PEM by at least 3-fold, but by as many as 5-fold in some regions of the brain. Although most PEM are likely a subset of CAM, there may be some distinguishing features of PEM. In the adult brain over 80% of juxtavascular microglia remain adjacent to the vasculature over a six week period when imaged through cranial windows^13^. This finding was supported by Bisht et al., 2021 who identified ∼70% of CAM were stable over a four-week imaging period^15^. These findings differ from our data where we identified that less than half (44%) of the PEM we tracked over 28 days remained adjacent to pericytes throughout the whole imaging period. These findings suggest PEM may be a uniquely motile subset of CAM.

We observed microglia in contact with vessels and pericytes at areas lacking astrocyte endfeet, corroborating studies showing microglia contribute to the glia limitans^8,37^, and indicating communication could occur between pericytes and microglia without the structural interference of astrocyte endfeet. However, PEM do appear separated from pericytes by the basement membrane. This is akin to the relationship between pericytes and endothelial cells, whereby pericytes and endothelial cells are almost continuously separated by the basement membrane^38^. There are holes in the membrane that allow pericytes and endothelial cells to form connections through peg-and-socket type structures, within which junction proteins allow for the exchange of molecules between the cells^14,39^. It is not clear if such a relationship could exist between microglia and pericytes, but it will be important to determine this through detailed ultrastructural studies.

A number of signalling pathways have been described to connect pericytes and parenchymal cells, particularly astrocytes and neurons^40^. One candidate pathway that may play a role in the association and function of microglia at blood vessels in the CNS is the fractalkine (CX_3_CL1)/CX_3_CR1 pathway. A deficiency in the fractalkine receptor CX_3_CR1 delays the association of microglia with blood vessels in the developing brain^13^. Furthermore, in the brain and in retinal explants, the addition of CX_3_CL1 causes capillary constriction at regions of microglia contact^35^. The precise cellular source of CX_3_CL1 that could reproduce these effects *in vivo* is not yet clear, but capillary pericytes do have the capacity to release it^36^. In this study, we found no change in the number of pericyte-microglial associations in CX_3_CR1 KO mice, suggesting the CX_3_CL1/CX_3_CR1 axis may not be a major pathway for PEM recruitment or maintenance in the healthy adult mouse brain. This does not rule out a role for the CX_3_CL1/CX_3_CR1 axis being involved in microglia-pericyte signalling in disease conditions, especially considering pericytes increase the release of CX_3_CL1 in response to stimuli^36^.

There are many possible functions for the association between microglia and pericytes. One logical hypothesis is that PEM are involved in regulating capillary tone indirectly via pericytes. Pericytes are well established to control cerebral blood flow^20,41-45^, and signals received from other cells such as astrocytes have been shown to modulate capillary tone via pericytes^43^. We observed an increase in vessel width underneath both pericytes and CAM in the basal state and this increase was also observed when pericytes had a PEM, suggesting that pericytes and their PEM may modulate blood flow. However, further studies are needed, particularly assessing evoked capillary responses at sites of PEM and CAM, or in disease states to confirm a specific effect of this interaction on blood flow modulation.

Microglia migrate to vessels in response to various injurious stimuli including stroke^46^, traumatic brain injury^12^, systemic inflammation^14^, laser-induced vascular injury^47^, and experimental autoimmune encephalomyelitis^48,49^. In one study microglia were found to cluster around pericytes in a model of epilepsy^50^. Here, we observed the opposite phenomenon in the SFG of the AD brain, with a reduction in the proportion of microglia that were both CAM and PEM (Fig. 7). Our data supports an earlier study which found reduced microglia coverage within the neurovascular unit in the hippocampus and medial frontal gyrus of AD patients^30^. It is possible microglia are being recruited to scavenge excess Aβ that is being deposited in the extracellular space. Microglia have long been known to cluster around amyloid plaques and phagocytose Aβ^51,52^. A result of microglia recruitment to Aβ plaques could be the loss of their physiological function at blood vessels and pericytes, which may contribute to functional decline in AD. When we stratified our AD cohort by *APOE* status, there were significantly fewer PEM in people with *APOE* ε3/ε4 compared to *APOE* ε3/ε3, suggesting *APOE4* may exacerbate PEM loss in AD. The mechanism causing this could be related to the reduced migratory potential of microglia expressing *APOE4*^53^. Pericytes are also affected by *APOE4*, with a reduced capacity to induce extracellular matrix production and barrier formation *in vitro* compared to *APOE3* pericytes^54^.

**Figure 7.**
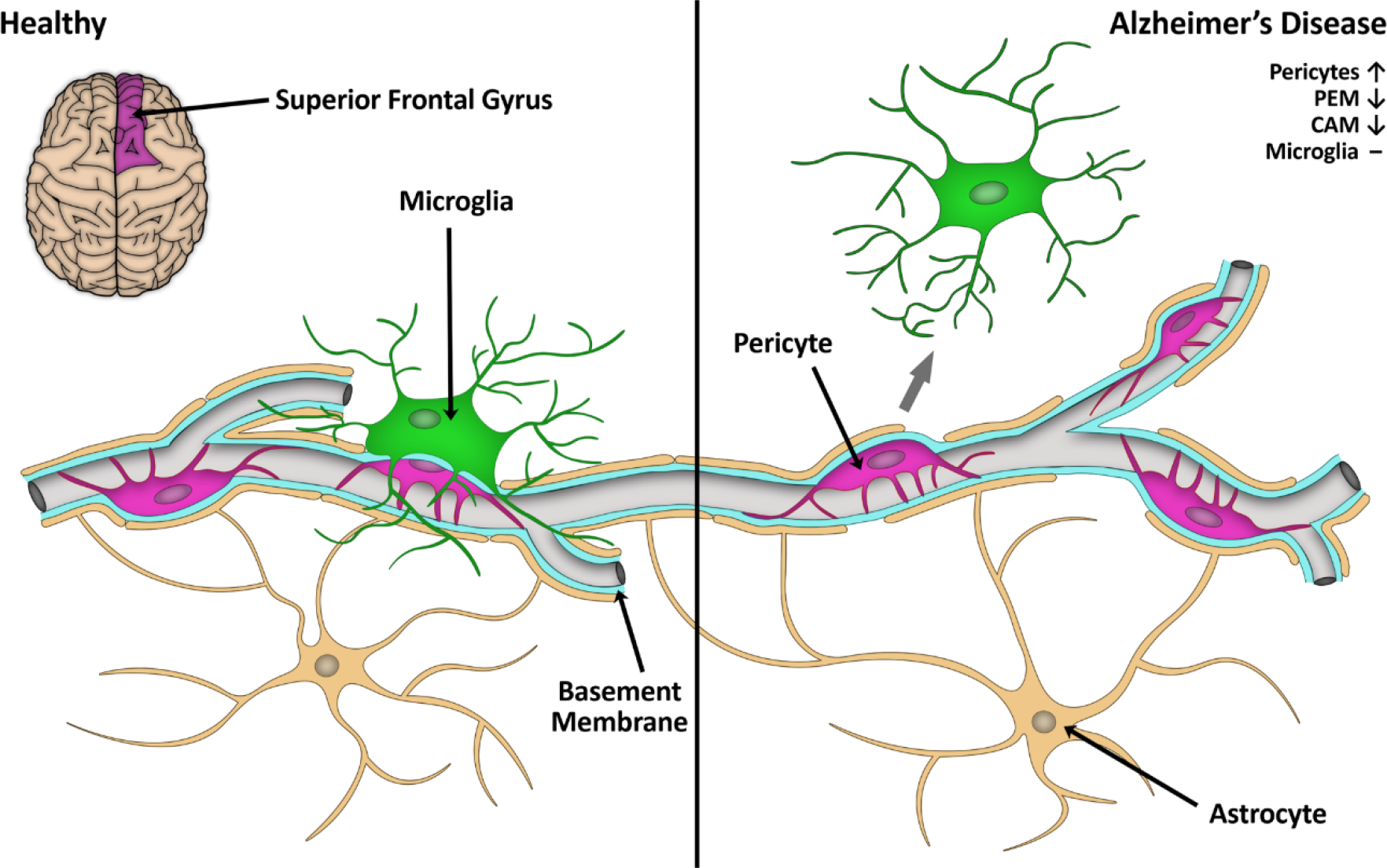
Schematic summary of pericyte and microglia associations in healthy control and AD. In this study, we found that pericyte density was increased, but the proportion of microglia that were PEM and CAM were reduced in the SFG of human AD brains compared to controls. This schematic also illustrates our finding that PEM can be found adjacent to pericytes lacking AQP4 positive astrocyte endfeet coverage. It is unknown if astrocyte endfeet coverage is altered in the SFG of AD, but endfeet coverage or function is hypothetically reduced, as suggested by others^68,69^.

Our finding that pericyte number was greater in the SFG in AD compared to controls is contrary to other studies where a decline in the number of pericytes was observed in the frontal cortex and hippocampus in human AD^28^, with an *APOE4* genotype appearing to exacerbate this pericyte loss^29^. Other studies have similarly found decreased pericyte coverage in the hippocampus and middle frontal gyrus as Braak stage increases^30,55^. However, two studies have recently challenged the notion that pericyte death is a general feature of AD pathology. One study found pericyte density was 28% greater in the middle frontal gyrus of AD cases, compared to age-matched controls^56^. A second study found a significant increase in the density of pericytes per length of vessels in the medial/dorso-lateral frontal cortex of AD cases compared to controls^57^. Our results corroborate these studies by showing higher pericyte numbers in AD compared to controls in an adjacent region, the SFG, and therefore provide further evidence that pericyte death is not necessarily a feature within every brain region in AD.

In summary, we have defined a novel subpopulation of microglia that reside directly adjacent to pericytes in the healthy brain, and we have termed these PEM. PEM are consistent in their distribution throughout brain and spinal cord regions in the healthy young adult mouse brain, and could serve a number of important functional roles including the modulation of blood flow. However, in clinical AD that proportion of microglia that are PEM is reduced in the SFG, which may have implications for the physiological function of microglia at the vasculature. Our work provides a platform to begin understanding the functions and signalling mechanisms controlling the communication between pericytes and microglia, and how a breakdown in the associations may contribute to the development of AD and other neurological diseases.

## Materials and methods

### Animals

All animal procedures were approved by the Animal Ethics Committee, University of Tasmania (A18608 and A23735) and conformed with the Australian National Health and Medical Research Council (NHMRC) Code of Practice for the Care and Use of Animals for Scientific Purposes, 2013 (8th Ed.). All results are reported in accordance with the ARRIVE guidelines^58^. Hemizygote NG2DsRed transgenic mice (The Jackson Laboratory stock #008241) were backcrossed onto a C57BL/6J background and crossbred with CX_3_CR1^GFP/GFP^ transgenic mice (The Jackson Laboratory stock #005582, C57BL/6J background) to produce NG2DsRed x CX_3_CR1^+/GFP^ or NG2DsRed x CX_3_CR1^GFP/GFP^ mice. A total of 29 mice were used throughout the study. Mice were group housed in Optimouse caging on a 12/12 h light/dark cycle with *ad libitum* access to food and water, with acclimation for ≥7 days.

### Tissue collection, processing and image collection

#### Tissue Collection

12-week-old NG2DsRed x CX_3_CR1^+/GFP^ or NG2DsRed x CX_3_CR1^GFP/GFP^ mice were given a lethal intraperitoneal injection of pentobarbitone (300 mg/kg) and immediately transcardially perfused with heparinised PBS followed by 4% paraformaldehyde (PFA; pH 7.4). Whole brains were prepared for cryosectioning as previously described^59^.

#### Blood vessel labelling and immunohistochemistry for astrocyte endfeet

For isolectin labelling one section per brain was slide mounted, air dried and permeabilised in PBS with 1% Triton X-100 for 20 min at RT. Sections were incubated overnight at 4°C with isolectin GS-IB4 (1:100; Invitrogen, Cat# I32450) in PBS with 1% Triton X-100. Sections were washed with PBS (3 × 5 min) and treated with Trueblack (Biotium, Cat# 23007) according to the manufacturer’s instructions. Sections were washed in PBS (3 × 5 min), rinsed in distilled water, briefly airdried, and coverslipped with Prolong Gold with DAPI (Life Technologies, Cat# P36935).

Sections for AQP4 labelling were slide mounted, permeabilised in PBS with 0.3% Triton X-100 for 40 min, washed in PBS (3 × 5min) and incubated with serum free protein block (Dako, Cat# X090930-2) for 1h at RT. Sections were incubated overnight at 4°C with guinea pig-anti-AQP4 antibody (1:250; Synaptic Systems, Cat# 429004) in antibody diluent (Dako, Cat# S080983-2). Sections were rinsed with PBS (3x 5 min) and incubated in donkey-anti-guinea pig 647 (1:1000; ThermoFisher, Cat# A21450) in antibody diluent for 2h at RT. Sections were washed in PBS (3 × 5 min), treated with Trueblack, washed in PBS (3 × 5 min), rinsed in distilled water, briefly airdried and cover slipped with Prolong Gold with DAPI.

### Imaging of CAM/PEM in the mouse brain

DAPI ± isolectin stained NG2DsRed x CX_3_CR1^+/GFP^ and NG2DsRed x CX_3_CR1^GFP/GFP^ tissue was imaged using a VS120 Virtual Slide System (Olympus, Japan). Sections used for CAM/PEM analysis were imaged at 40x magnification across five focal planes, spaced 2 μm apart, using the extended focal imaging setting. Optimum exposure times for DAPI (Ex: 388 nm; Em: 448 nm), DsRed (Ex: 576 nm; Em: 625 nm), GFP (Ex: 494 nm; Em: 530 nm) and isolectin (Ex: 640 nm; Em: 690 nm) were kept consistent for all images in each cohort.

Confocal stacks of isolectin and AQP4 labelled NG2DsRed x CX_3_CR1^+/GFP^ tissue were spaced at 1.0 or 0.5 μm increments, using an inverted Ti Eclipse microscope (Nikon, Japan) equipped with a CSU-X1 spinning disk scanner (Yokogawa Electric Corporation, Japan). Images were acquired with a 100x (1.40) oil objective (Nikon, Japan) with filters for DAPI (405/445), FITC (488/525), TRITC (561/615), and CY5 (640/705) and analysed using NIS-Elements AR 5.02.00 (Nikon, Japan). Images were exported as OME TIFFs and NIS-Elements AR 5.02.00 was used to create 3D reconstructions.

### Cranial widow implantation and *in vivo* two-photon imaging

Cranial windows were implanted in nine 12-week-old NG2DsRed x CX_3_CR1^+/GFP^ mice as described previously^60^, with modification; windows were implanted without a titanium bar glued opposite the window and were placed over the primary somatosensory cortex, specifically over the upper, lower and trunk domain areas. *In vivo* two-photon laser scanning microscopy (2PLSM) was performed using the same microscope and software as previously described^60^ and was performed during the mouse light cycle. To image blood vessels, mice were intravenously (via the tail vein) administered 2% w/v FITC-dextran in saline (70,000 Da; Sigma-Aldrich, USA) 10 minutes prior to imaging. Mice were anesthetised with isoflurane in a chamber, then transferred to a stereotaxic frame, where isoflurane was delivered through a facemask at 2-3% concentration in oxygen, as required, to maintain anaesthesia^60^. An EC Plan-Neofluar 20X/0.1 water immersion objective (Nikon, USA), illuminated by white light, was used to identify large blood vessels in the somatosensory cortex (layers II/III) that were subsequently used as landmarks for regions of interest (ROIs). The resultant brightfield image was captured using XCAP Image Processing Software (EPIX Incorporated, USA). Once ROIs were identified for imaging, z-plane coordinates were saved. The same z-plane coordinate was initiated using the 2PLSM, and a higher-resolution ROI captured. Femto second-pulsed infrared excitation was from a mode-locked Ti-sapphire laser tuned to 910 nm and equipped with group velocity dispersion compensation (Mai Tai DeepSee; Spectra-Physics, Australia). Power delivered to the back aperture was 40-100mW (14-17% power), depending on depth. This approach allowed the visualisation of NG2DsRed, CX_3_CR1^+/GFP^ and fluorescein isothiocyanate (FITC)-dextran signals, simultaneously. All ROIs were imaged with a zoom of 2.0 (210 μm x 210 μm field of view) with 1 μm z-step intervals, beginning from a depth of 20 μm below the brain’s surface down to 130 μm.

### Quantification of pericytes, microglia, CAM and PEM in a human Alzheimer’s disease cohort

#### Human cohort details

Research using human tissue was approved by the University of Tasmania’s Human Medical Research Ethics Committee (H20078) in accordance with the NHMRC National Statement on Ethical Conduct in Human Research (2018). Human brain tissue was obtained from the Banner Sun Health Research Institute’s Brain and Body Donation Program. All subjects, or their legal representatives, signed an Institutional Review Board-approved informed consent form for brain donation (for more information see^61^). Briefly, human brain tissue was fixed with 4% formaldehyde, cryoprotected in 2% dimethyl sulfoxide/20% glycerol, sectioned at 40 μm on a sledge-type microtome and free-floating sections stored in PBS with 0.02% sodium azide at 4°C.

Sections from the superior frontal gyrus (SFG) of 12 control cases and 11 AD cases were used in the current study (Table 1). Cases were considered to have AD if, at a minimum, they were defined as intermediate or high on the NIA-Reagan criteria^61,62^. A case was considered a control if the person did not have dementia or parkinsonism during life and was without any major neuropathological diagnosis. The SFG was selected for the current analysis as it is an area susceptible to the development of amyloid plaques and tau tangles in AD^63,64^.

**Table 1.** Demographics of human brain tissue cases.

| *Demographics* | *Control (n=12)* | *AD (n=11)* |
| --- | --- | --- |
| *Number of cases by sex: male/female (percentage of cases)* | 5 (42%) / 7 (58%) | 5 (45%) / 6 (55%) |
| *Mean age at death: years (range)* | 83.8 (53-95) | 80.2 (61-89) |
| *Mean post-mortem interval: hours (range)* | 2.88 (2-3.95) | 2.73 (1.75-5) |
| *Number of cases with each APOE Genotype* |  |  |
| *ε2/ε2* | 1 | - |
| *ε3/ε3* | 10 | 5 |
| *ε3/ε4* | - | 6 |
| *ε4/ε4* | 1 | - |

#### Labelling and imaging of pericytes, microglia and blood vessels in human brain sections

Free-floating human SFG sections were permeabilised in PBS with 0.3% Triton X-100 for 1h at RT, incubated in citric acid antigen retrieval buffer (pH 4.5) overnight at 4°C, and then microwaved (650W, 30s) in 10mL of fresh citric acid antigen retrieval buffer. Sections were cooled to RT for 30min, washed in PBS with 0.3% Triton X-100 (3 × 5 min), and incubated in serum free blocking solution (Dako, Cat# X090930-2) for 1h at RT. Sections were incubated at 4°C for three nights with rabbit-anti-PDGFRβ (1:100; Thermofisher, Cat# ab32570), guinea pig-anti-IBA1 (1:500; Synaptic Systems, Cat# 234004) and UEA-1 594 (1:1000; Vector Laboratories, Cat# DL-1067), in antibody diluent (Dako, Cat #S0809). Sections were rinsed in PBS with 0.3% Triton X-100 (3 × 10 min) and then incubated with donkey-anti-guinea pig 647 (1:1000; ThermoFisher Cat# A21450) and donkey-anti-rabbit 488 (1:1000; ThermoFisher Cat# A21206) overnight at 4°C. Sections were washed in PBS with 0.3% Triton X-100 (3 × 10 min), incubated in PBS containing DAPI (1:5000, Invitrogen, Cat#D3571, 1x 10 min), slide mounted, briefly air dried and incubated in Trueblack, according to the manufacturer’s instructions (Biotium, Cat# 23007). Sections were rinsed with distilled water and cover slipped with fluorescent mounting medium (Fischer Scientific, Cat# TA-030-FM). Human tissue was imaged with both a VS120 Virtual Slide System and confocal microscope using the same protocols described above for isolectin stained NG2DsRed x CX_3_CR1^+/GFP^ mouse brain sections. The researcher performing immunohistochemistry was blinded to case type throughout experimental procedures and analysis.

### Image analysis: NG2DsRed x CX_3_CR1^+/GFP or GFP/GFP^ mouse brains

#### CAM and PEM prevalence in the somatosensory cortex

CAM and PEM prevalence was manually quantified in isolectin labelled NG2DsRed x CX_3_CR1^+/GFP^ tissue using QuPath 0.3.2^65^. Boxed annotations (800 × 600 μm) were placed in the somatosensory cortex of both hemispheres. All in focus microglia and pericytes containing a DAPI-positive nucleus were counted. Cells were not counted if their nuclei touched the edge of the box. DsRed-positive pericytes were only counted if they were associated with a vessel (specifically vessels <10 μm in diameter). Microglia were counted as a CAM if the centre of their nuclei were <10 μm from the centre of the closest isolectin-positive blood vessel (specifically vessels <10 μm in diameter), similar to previous approaches^13,15^. Microglia were counted as PEM if the centre point of the microglia nucleus was <10 μm from the centre point of a pericyte nucleus (specifically pericytes on vessels <10 μm in diameter). To determine if microglia and pericyte associations occur more than expected by chance Monte-Carlo simulations were performed, as described in Supplementary Methods.

Microglia, pericytes, CAM and PEM were also analysed in isolectin-stained brain sections from NG2DsRed x CX_3_CR1^GFP/GFP^ mice and compared to a different cohort of 12-week-old NG2DsRed x CX_3_CR1^+/GFP^ mice using the same manual protocol above, with modification. The same somatosensory regions were traced and a 400 × 400 μm grid was placed within the traced areas. Microglia and pericytes were counted in the first complete 400 × 400 μm square within the annotation, with every second complete square thereafter counted to avoid sampling bias, until six full boxes per hemisphere were counted (12 counted in total per brain). Any incomplete squares or squares containing artefacts (e.g. tissue rips, folds or other artefacts that affected cell counting) were skipped. Cells contacting the edges of the squares were only counted if their nuclei were touching the top, or left-hand margins of the box, to avoid double counting. The researcher performing the analysis was blinded to genotype throughout quantification.

### Image Analysis: *in vivo* NG2DsRed x CX_3_CR1^+/GFP^ two-photon derived images

#### Quantification of CAM and PEM

Using FIJI-ImageJ (NIH, USA)^66^, images were automatically adjusted for brightness/contrast. The plugin ‘Cell Counter’ was used to manually annotate the pericytes and microglia in each ROI. Microglial GFP signal and FITC-dextran vessel lumen signal was differentiated by the higher intensity of the GFP compared to the FITC and the presence of dark lines indicating red blood cells within the vessel lumen. The distances between pericytes and microglia were calculated based on x, y, and z co-ordinates for the centre point of each cell soma, using FIJI-ImageJ.

#### Quantification of pericyte, CAM and PEM vascular tree location

Using images obtained on imaging day -1 (with FITC-dextran, see Fig. 2), the position of vessels within the vascular tree was able to be determined through z-stacks of 10 ROIs across five mice, as previously illustrated^20^. Penetrating arterioles (0 order) were identified by their large luminal size and presence of NG2DsRed-positive rings of vascular smooth muscle cells (VSMCs). Branching higher order capillaries (1st-2nd order) were identified by their smaller vessel width and presence of transitional ensheathing pericytes. Mid-order (3rd-4th) and lower order capillaries (≥5th order) were identified by their small vessel width and presence of thin-strand pericytes. Ascending venules (8th order) were identified by their larger luminal size and mesh pericyte processes with a distinct lack of ringed VSMCs. See Fig. 2 for an example of a ROI that encompassed an entire vascular tree. We then quantified the position of pericytes, CAM and PEM at different levels of the vascular tree.

#### Tracking microglia proximity to pericytes over time

The x, y, and z co-ordinates of all microglia and pericytes within a ROI was used to determine the relative distance between the cells on each day of imaging. On day 0, 32 pericytes from six mice were identified to have a PEM. These 32 pericytes with a PEM on day 0, were re-imaged on days 4, 7 and 28 to determine if microglial associations with pericytes are stable or dynamic. The total number of pericytes and pericytes with a PEM in each ROI was also quantified on each imaging day, to calculate the percentage of pericytes with a PEM on days 0, 4, 7 and 28.

#### Determining vessel width at locations of CAM, PEM, and pericytes along capillaries

Using the line tool in FIJI-ImageJ, images taken on day -1 (when FITC-dextran was administered to visualise vessels) were manually annotated to measure vessel width of capillaries at four specific locations: vessel only (VO), containing no pericyte or microglia cell bodies; pericyte on vessel (P), where a pericyte was present alone on a vessel; CAM, where a microglia alone was associated with a vessel; and PEM, where a microglia was associated with a pericyte on a vessel (Fig. 4F). All measurements were made on vessel segments without branchpoints. For VO, a line was drawn across the vessel. For pericytes, CAM and PEM, a line was drawn across the vessel at the centre of the pericyte (for pericytes and PEM) or microglial (for CAM) cell bodies.

### Image Analysis: pericytes, microglia, CAM and PEM in human brain sections

Pericytes and microglia were manually counted using QuPath 3.0. ROI annotations were created by tracing the entire grey matter in each section, and then shrinking the trace by 50 μm to remove edge artefacts. Other obvious artefacts (e.g. tissue rips, folds, etc) were manually removed from the annotations. A 500 × 500 μm grid was placed within each annotation. Microglia and pericytes were counted in the first complete 500 × 500 μm square within the annotation, with every fourth complete square counted thereafter to avoid sampling bias, until twenty full squares were counted. Any incomplete squares or squares containing artefacts (e.g. tissue rips, folds or other artefacts that affected cell counting) were skipped. Cells contacting the edges of the squares were only counted if their nuclei were touching the top, or left-hand margins of the box, to avoid double counting. Cells were only counted if there was a clear DAPI-positive nucleus present. The researcher performing the analysis was blinded to genotype throughout quantification.

### Statistical analysis

Data were processed in Microsoft Excel and all statistical analysis was performed using GraphPad Prism 9.3.1 (GraphPad, USA). Prior to statistical analysis, all data underwent outlier testing using a ROUT test (Q = 1%). Data were tested for normality using the D’Agostino & Pearson test, or the Shapiro-Wilk test if n numbers were too small for the D’Agostino & Pearson. For unpaired two group comparisons, means were compared using an unpaired t-test, if data were normally distributed, but a Welch’s correction was applied if variance was significantly different as tested by an F test. If data were not normally distributed in any group, a Mann-Whitney test was employed. For paired two group comparisons, a paired t-test was applied if data were normally distributed, and a Wilcoxon test applied if data were not normally distributed in any group. For comparisons with more than two groups, if data were normally distributed a repeated measures one-way ANOVAs with a Geisser-Greenhouse correction was applied to account for variations in sphericity and Tukey’s post-hoc test employed for multiple comparisons. If repeated measures data were not normally distributed a Friedman test followed by a Dunn’s post-hoc test was applied. A *p* < 0.05 was considered statistically significant. Statistical tests used for each analysis are reported in figure legends. All data will be made available upon request.

## Author contributions

GPM: conceptualization, data curation, validation, formal analysis, investigation, methodology, visualization, software, project administration, writing – original draft, review and editing; CGF: conceptualization, data curation, validation, formal analysis, investigation, methodology, visualization, software, writing – original draft, review and editing; JoMC: data curation, methodology, resources, software, writing – review & editing; JeMC: investigation, resources, methodology, writing – review & editing; JaMC: methodology, investigation, writing – review & editing; LSB: methodology, software, validation, writing – review & editing; DWH: conceptualisation, supervision, writing – review & editing; GCD: conceptualisation, supervision, funding acquisition, writing – review & editing; AJC: conceptualisation, supervision, methodology, writing – review & editing; AEK: funding acquisition, resources, writing – review & editing, visualization, supervision; JMZ: conceptualization, methodology, investigation, resources, writing – original draft, review and editing, supervision; BAS: conceptualization, methodology, formal analysis, resources, writing – original draft, review and editing, visualization, supervision, project administration, funding acquisition.

## Acknowledgements

We thank Dr. Bill Bennett for technical assistance with two-photon microscopy imaging and Dr. Robert Gasperini for assistance with confocal microscopy throughout the project. We thank Dr. Lila Landowski for their valuable discussions and mentorship of the junior researchers associated with this project. We thank Dr. Dino Premilovac for valuable discussions about the study. We thank Prof. James Vickers for providing access to the human tissue cohort used in this study and we thank the Banner Sun Health Research Institute’s Brain and Body Donation Program for generating the human tissue cohort used in this study.

## Abbreviations

2PLSM: two-photon laser scanning microscopy
Aβ: Amyloid beta
AD: Alzheimer’s disease
ANOVA: Analysis of variance
APOE: Apolipoprotein E
ARRIVE: Animal Research: Reporting of In Vivo Experiments
AQP4: aquaporin-4
CAM: Capillary-associated microglia
Cy5: Cyanine 5
CX_3_CL1: C-X3-C Motif Chemokine Ligand 1
CX_3_CR1: C-X3-C chemokine receptor 1
DAPI: 4′,6-diamidino-2-phenylindole
FITC: Fluorescein isothiocyanate
GFP: Green Fluorescent Protein
IBA1: Ionized calcium binding adaptor molecule 1
KO: Knockout
NHMRC: National Health and Medical Research Council
NIA: National Institute on Aging
NIH: National Institutes of Health
OME TIFF: Open Microscopy Environment Tag Image File Format
PBS: phosphate buffered saline
PDGFRβ: Platelet-derived growth factor receptor beta
PEM: Pericyte-associated microglia
PFA: Paraformaldehyde
ROI/s: Region/s of interest
ROUT: Robust regression and outlier removal
RT: Room temperature
SFG: Superior frontal gyrus
TRITC: Tetramethylrhodamine
UEA-1: Ulex Europaeus Agglutinin I

## References

1 Werneburg, S., Feinberg, P. A., Johnson, K. M. & Schafer, D. P. A microglia-cytokine axis to modulate synaptic connectivity and function. Curr Opin Neurobiol 47, 138–145, doi:10.1016/j.conb.2017.10.002 (2017).

2 Wright-Jin, E. C. & Gutmann, D. H. Microglia as Dynamic Cellular Mediators of Brain Function. Trends Mol Med 25, 967–979, doi:10.1016/j.molmed.2019.08.013 (2019).

3 Prinz, M., Masuda, T., Wheeler, M. A. & Quintana, F. J. Microglia and Central Nervous System–Associated Macrophages—From Origin to Disease Modulation. Annual Review of Immunology 39, 251–277, doi:10.1146/annurev-immunol-093019-110159 (2021).

4 Cammermeyer, J. The hypependymal microglia cell. Zeitschrift für Anatomie und Entwicklungsgeschichte 124, 543–561 (1965).

5 Rezaie, P., Cairns, N. J. & Male, D. K. Expression of adhesion molecules on human fetal cerebral vessels: relationship to microglial colonisation during development. Brain Res Dev Brain Res 104, 175–189, doi:10.1016/s0165-3806(97)00153-3 (1997).

6 Cuadros, M. A., Martin, C., Coltey, P., Almendros, A. & Navascués, J. First appearance, distribution, and origin of macrophages in the early development of the avian central nervous system. Journal of Comparative Neurology 330, 113–129, doi:https://doi.org/10.1002/cne.903300110 (1993).

7 Barón, M. & Gallego, A. The relation of the microglia with the pericytes in the cat cerebral cortex. Z Zellforsch Mikrosk Anat 128, 42–57, doi:10.1007/bf00306887 (1972).

8 Lassmann, H., Zimprich, F., Vass, K. & Hickey, W. F. Microglial cells are a component of the perivascular glia limitans. J Neurosci Res 28, 236–243, doi:https://doi.org/10.1002/jnr.490280211 (1991).

9 Oemichen, M. Mononuclear phagocytes in the central nervous system. Origin, mode of distribution, and function of progressive microglia, perivascular cells of intracerebral vessels, free subarachnoidal cells, and epiplexus cells. Schriftenr Neurol 21, I-x, 1-167 (1978).

10 Rio Hortega, P. d. La microglia y su transformacion en celulas en bastoncito yen cuerpos granuloadiposos. Trab. Lab. Invest. Biol. (1920).

11 Graeber, M. B. & Streit, W. J. Perivascular microglia defined. Trends in Neurosciences 13, 366, doi:https://doi.org/10.1016/0166-2236(90)90020-B (1990).

12 Grossmann, R., Stence, N., Carr, J., Fuller, L., Waite, M. & Dailey, M. E. Juxtavascular microglia migrate along brain microvessels following activation during early postnatal development. Glia 37, 229–240 (2002).

13 Mondo, E., Becker, S. C., Kautzman, A. G., Schifferer, M., Baer, C. E., Chen, J., Huang, E. J., Simons, M. & Schafer, D. P. A Developmental Analysis of Juxtavascular Microglia Dynamics and Interactions with the Vasculature. The Journal of Neuroscience 40, 6503–6521, doi:10.1523/jneurosci.3006-19.2020 (2020).

14 Haruwaka, K., Ikegami, A., Tachibana, Y., Ohno, N., Konishi, H., Hashimoto, A., Matsumoto, M., Kato, D., Ono, R., Kiyama, H., Moorhouse, A. J., Nabekura, J. & Wake, H. Dual microglia effects on blood brain barrier permeability induced by systemic inflammation. Nature communications 10, 5816–5816, doi:10.1038/s41467-019-13812-z (2019).

15 Bisht, K., Okojie, K. A., Sharma, K., Lentferink, D. H., Sun, Y.-Y., Chen, H.-R., Uweru, J. O., Amancherla, S., Calcuttawala, Z., Campos-Salazar, A. B., Corliss, B., Jabbour, L., Benderoth, J., Friestad, B., Mills, W. A., Isakson, B. E., Tremblay, M.-é., Kuan, C.-Y. & Eyo, U. B. Capillary-associated microglia regulate vascular structure and function through PANX1-P2RY12 coupling in mice. Nature Communications 12, 5289, doi:10.1038/s41467-021-25590-8 (2021).

16 Császár, E., Lénárt, N., Cserép, C., Környei, Z., Fekete, R., Pósfai, B., Balázsfi, D., Hangya, B., Schwarcz, A. D., Szabadits, E., Szöllősi, D., Szigeti, K., Máthé, D., West, B. L., Sviatkó, K., Brás, A. R., Mariani, J.-C., Kliewer, A., Lenkei, Z., Hricisák, L., Benyó, Z., Baranyi, M., Sperlágh, B., Menyhárt, Á., Farkas, E. & Dénes, Á. Microglia modulate blood flow, neurovascular coupling, and hypoperfusion via purinergic actions. Journal of Experimental Medicine 219, doi:10.1084/jem.20211071 (2022).

17 Kisler, K., Nikolakopoulou, A. M. & Zlokovic, B. V. Microglia have a grip on brain microvasculature. Nat Commun 12, 5290, doi:10.1038/s41467-021-25595-3 (2021).

18 Brown, L. S., Foster, C. G., Courtney, J.-M., King, N. E., Howells, D. W. & Sutherland, B. A. Pericytes and Neurovascular Function in the Healthy and Diseased Brain. Front Cell Neurosci 13, 282–282, doi:10.3389/fncel.2019.00282 (2019).

19 Sutherland, B. A. In Biology of Pericytes – Recent Advances (ed Alexander Birbrair) 39-74 (Springer International Publishing, 2021).

20 Hall, C. N., Reynell, C., Gesslein, B., Hamilton, N. B., Mishra, A., Sutherland, B. A., O’Farrell, F. M., Buchan, A. M., Lauritzen, M. & Attwell, D. Capillary pericytes regulate cerebral blood flow in health and disease. Nature 508, 55–60, doi:10.1038/nature13165 (2014).

21 Risau, W. & Wolburg, H. Development of the blood-brain barrier. Trends in Neurosciences 13, 174–178, doi:https://doi.org/10.1016/0166-2236(90)90043-A (1990).

22 Sakuma, R., Kawahara, M., Nakano-Doi, A., Takahashi, A., Tanaka, Y., Narita, A., Kuwahara-Otani, S., Hayakawa, T., Yagi, H., Matsuyama, T. & Nakagomi, T. Brain pericytes serve as microglia-generating multipotent vascular stem cells following ischemic stroke. J Neuroinflammation 13, 57–57, doi:10.1186/s12974-016-0523-9 (2016).

23 Özen, I., Deierborg, T., Miharada, K., Padel, T., Englund, E., Genové, G. & Paul, G. Brain pericytes acquire a microglial phenotype after stroke. Acta Neuropathol 128, 381–396, doi:10.1007/s00401-014-1295-x (2014).

24 Toledo, J. B., Arnold, S. E., Raible, K., Brettschneider, J., Xie, S. X., Grossman, M., Monsell, S. E., Kukull, W. A. & Trojanowski, J. Q. Contribution of cerebrovascular disease in autopsy confirmed neurodegenerative disease cases in the National Alzheimer’s Coordinating Centre. Brain 136, 2697–2706, doi:10.1093/brain/awt188 (2013).

25 Steinman, J., Sun, H.-S. & Feng, Z.-P. Microvascular Alterations in Alzheimer’s Disease. Front Cell Neurosci 14, doi:10.3389/fncel.2020.618986 (2021).

26 Nortley, R., Korte, N., Izquierdo, P., Hirunpattarasilp, C., Mishra, A., Jaunmuktane, Z., Kyrargyri, V., Pfeiffer, T., Khennouf, L., Madry, C., Gong, H., Richard-Loendt, A., Huang, W., Saito, T., Saido, T. C., Brandner, S., Sethi, H. & Attwell, D. Amyloid β oligomers constrict human capillaries in Alzheimer’s disease via signaling to pericytes. Science 365, eaav9518, doi:10.1126/science.aav9518 (2019).

27 Sagare, A. P., Bell, R. D., Zhao, Z., Ma, Q., Winkler, E. A., Ramanathan, A. & Zlokovic, B. V. Pericyte loss influences Alzheimer-like neurodegeneration in mice. Nat Commun 4, 2932, doi:10.1038/ncomms3932 (2013).

28 Sengillo, J. D., Winkler, E. A., Walker, C. T., Sullivan, J. S., Johnson, M. & Zlokovic, B. V. Deficiency in mural vascular cells coincides with blood-brain barrier disruption in Alzheimer’s disease. Brain Pathol 23, 303–310, doi:10.1111/bpa.12004 (2013).

29 Halliday, M. R., Rege, S. V., Ma, Q., Zhao, Z., Miller, C. A., Winkler, E. A. & Zlokovic, B. V. Accelerated pericyte degeneration and blood-brain barrier breakdown in apolipoprotein E4 carriers with Alzheimer’s disease. J Cereb Blood Flow Metab 36, 216–227, doi:10.1038/jcbfm.2015.44 (2016).

30 Kirabali, T., Rust, R., Rigotti, S., Siccoli, A., Nitsch, R. M. & Kulic, L. Distinct changes in all major components of the neurovascular unit across different neuropathological stages of Alzheimer’s disease. Brain Pathology 30, 1056–1070, doi:https://doi.org/10.1111/bpa.12895 (2020).

31 Jonsson, T., Stefansson, H., Steinberg, S., Jonsdottir, I., Jonsson, P. V., Snaedal, J., Bjornsson, S., Huttenlocher, J., Levey, A. I., Lah, J. J., Rujescu, D., Hampel, H., Giegling, I., Andreassen, O. A., Engedal, K., Ulstein, I., Djurovic, S., Ibrahim-Verbaas, C., Hofman, A., Ikram, M. A., van Duijn, C. M., Thorsteinsdottir, U., Kong, A. & Stefansson, K. Variant of TREM2 Associated with the Risk of Alzheimer’s Disease. New England Journal of Medicine 368, 107–116, doi:10.1056/NEJMoa1211103 (2012).

32 Schwabe, T., Srinivasan, K. & Rhinn, H. Shifting paradigms: The central role of microglia in Alzheimer’s disease. Neurobiology of Disease 143, 104962, doi:https://doi.org/10.1016/j.nbd.2020.104962 (2020).

33 Hong, S., Beja-Glasser, V. F., Nfonoyim, B. M., Frouin, A., Li, S., Ramakrishnan, S., Merry, K. M., Shi, Q., Rosenthal, A., Barres, B. A., Lemere, C. A., Selkoe, D. J. & Stevens, B. Complement and microglia mediate early synapse loss in Alzheimer mouse models. Science 352, 712–716, doi:10.1126/science.aad8373 (2016).

34 Keren-Shaul, H., Spinrad, A., Weiner, A., Matcovitch-Natan, O., Dvir-Szternfeld, R., Ulland, T. K., David, E., Baruch, K., Lara-Astaiso, D., Toth, B., Itzkovitz, S., Colonna, M., Schwartz, M. & Amit, I. A Unique Microglia Type Associated with Restricting Development of Alzheimer’s Disease. Cell 169, 1276-1290.e1217, doi:10.1016/j.cell.2017.05.018 (2017).

35 Mills, S. A., Jobling, A. I., Dixon, M. A., Bui, B. V., Vessey, K. A., Phipps, J. A., Greferath, U., Venables, G., Wong, V. H. Y., Wong, C. H. Y., He, Z., Hui, F., Young, J. C., Tonc, J., Ivanova, E., Sagdullaev, B. T. & Fletcher, E. L. Fractalkine-induced microglial vasoregulation occurs within the retina and is altered early in diabetic retinopathy. Proceedings of the National Academy of Sciences 118, e2112561118, doi:10.1073/pnas.2112561118 (2021).

36 Smyth, L. C. D., Rustenhoven, J., Park, T. I. H., Schweder, P., Jansson, D., Heppner, P. A., O’Carroll, S. J., Mee, E. W., Faull, R. L. M., Curtis, M. & Dragunow, M. Unique and shared inflammatory profiles of human brain endothelia and pericytes. J Neuroinflammation 15, 138–138, doi:10.1186/s12974-018-1167-8 (2018).

37 Mathiisen, T. M., Lehre, K. P., Danbolt, N. C. & Ottersen, O. P. The perivascular astroglial sheath provides a complete covering of the brain microvessels: an electron microscopic 3D reconstruction. Glia 58, 1094–1103, doi:10.1002/glia.20990 (2010).

38 Armulik, A., Genové, G. & Betsholtz, C. Pericytes: Developmental, Physiological, and Pathological Perspectives, Problems, and Promises. Developmental Cell 21, 193–215, doi:https://doi.org/10.1016/j.devcel.2011.07.001 (2011).

39 ElAli, A., Thériault, P. & Rivest, S. The role of pericytes in neurovascular unit remodeling in brain disorders. Int J Mol Sci 15, 6453–6474, doi:10.3390/ijms15046453 (2014).

40 Sweeney, M. D., Ayyadurai, S. & Zlokovic, B. V. Pericytes of the neurovascular unit: key functions and signaling pathways. Nature Neuroscience 19, 771–783, doi:10.1038/nn.4288 (2016).

41 Sieczkiewicz, G. J. & Herman, I. M. TGF-beta 1 signaling controls retinal pericyte contractile protein expression. Microvasc Res 66, 190–196, doi:10.1016/s0026-2862(03)00055-4 (2003).

42 Yemisci, M., Gursoy-Ozdemir, Y., Vural, A., Can, A., Topalkara, K. & Dalkara, T. Pericyte contraction induced by oxidative-nitrative stress impairs capillary reflow despite successful opening of an occluded cerebral artery. Nat Med 15, 1031–1037, doi:10.1038/nm.2022 (2009).

43 Mishra, A., Reynolds, J. P., Chen, Y., Gourine, A. V., Rusakov, D. A. & Attwell, D. Astrocytes mediate neurovascular signaling to capillary pericytes but not to arterioles. Nat Neurosci 19, 1619–1627, doi:10.1038/nn.4428 (2016).

44 Neuhaus, A. A., Couch, Y., Sutherland, B. A. & Buchan, A. M. Novel method to study pericyte contractility and responses to ischaemia in vitro using electrical impedance. J Cereb Blood Flow Metab 37, 2013–2024, doi:10.1177/0271678x16659495 (2017).

45 Peppiatt, C. M., Howarth, C., Mobbs, P. & Attwell, D. Bidirectional control of CNS capillary diameter by pericytes. Nature 443, 700–704, doi:10.1038/nature05193 (2006).

46 Jolivel, V., Bicker, F., Binamé, F., Ploen, R., Keller, S., Gollan, R., Jurek, B., Birkenstock, J., Poisa-Beiro, L., Bruttger, J., Opitz, V., Thal, S. C., Waisman, A., Bäuerle, T., Schäfer, M. K., Zipp, F. & Schmidt, M. H. H. Perivascular microglia promote blood vessel disintegration in the ischemic penumbra. Acta Neuropathol 129, 279–295, doi:10.1007/s00401-014-1372-1 (2015).

47 Lou, N., Takano, T., Pei, Y., Xavier, A. L., Goldman, S. A. & Nedergaard, M. Purinergic receptor P2RY12-dependent microglial closure of the injured blood–brain barrier. Proceedings of the National Academy of Sciences 113, 1074–1079, doi:10.1073/pnas.1520398113 (2016).

48 Davalos, D., Kyu Ryu, J., Merlini, M., Baeten, K. M., Le Moan, N., Petersen, M. A., Deerinck, T. J., Smirnoff, D. S., Bedard, C., Hakozaki, H., Gonias Murray, S., Ling, J. B., Lassmann, H., Degen, J. L., Ellisman, M. H. & Akassoglou, K. Fibrinogen-induced perivascular microglial clustering is required for the development of axonal damage in neuroinflammation. Nat Commun 3, 1227, doi:10.1038/ncomms2230 (2012).

49 Joost, E., Jordão, M. J. C., Mages, B., Prinz, M., Bechmann, I. & Krueger, M. Microglia contribute to the glia limitans around arteries, capillaries and veins under physiological conditions, in a model of neuroinflammation and in human brain tissue. Brain Structure and Function 224, 1301–1314, doi:10.1007/s00429-019-01834-8 (2019).

50 Klement, W., Garbelli, R., Zub, E., Rossini, L., Tassi, L., Girard, B., Blaquiere, M., Bertaso, F., Perroy, J., de Bock, F. & Marchi, N. Seizure progression and inflammatory mediators promote pericytosis and pericyte-microglia clustering at the cerebrovasculature. Neurobiol Dis 113, 70–81, doi:10.1016/j.nbd.2018.02.002 (2018).

51 Paresce, D. M., Ghosh, R. N. & Maxfield, F. R. Microglial Cells Internalize Aggregates of the Alzheimer’s Disease Amyloid β-Protein Via a Scavenger Receptor. Neuron 17, 553–565, doi:https://doi.org/10.1016/S0896-6273(00)80187-7 (1996).

52 D’Andrea, M. R., Cole, G. M. & Ard, M. D. The microglial phagocytic role with specific plaque types in the Alzheimer disease brain. Neurobiology of Aging 25, 675–683, doi:https://doi.org/10.1016/j.neurobiolaging.2003.12.026 (2004).

53 Cudaback, E., Li, X., Montine, K. S., Montine, T. J. & Keene, C. D. Apolipoprotein E isoform-dependent microglia migration. The FASEB Journal 25, 2082–2091, doi:https://doi.org/10.1096/fj.10-176891 (2011).

54 Yamazaki, Y., Shinohara, M., Yamazaki, A., Ren, Y., Asmann, Y. W., Kanekiyo, T. & Bu, G. ApoE (Apolipoprotein E) in Brain Pericytes Regulates Endothelial Function in an Isoform-Dependent Manner by Modulating Basement Membrane Components. Arterioscler Thromb Vasc Biol 40, 128–144, doi:10.1161/atvbaha.119.313169 (2020).

55 Kirabali, T., Rigotti, S., Siccoli, A., Liebsch, F., Shobo, A., Hock, C., Nitsch, R. M., Multhaup, G. & Kulic, L. The amyloid-β degradation intermediate Aβ34 is pericyte-associated and reduced in brain capillaries of patients with Alzheimer’s disease. Acta Neuropathol Commun 7, 194–194, doi:10.1186/s40478-019-0846-8 (2019).

56 Fernandez-Klett, F., Brandt, L., Fernández-Zapata, C., Abuelnor, B., Middeldorp, J., Sluijs, J. A., Curtis, M., Faull, R., Harris, L. W., Bahn, S., Hol, E. M. & Priller, J. Denser brain capillary network with preserved pericytes in Alzheimer’s disease. Brain pathology (Zurich, Switzerland) 30, 1071–1086, doi:10.1111/bpa.12897 (2020).

57 Ding, R., Hase, Y., Burke, M., Foster, V., Stevenson, W., Polvikoski, T. & Kalaria, R. N. Loss with ageing but preservation of frontal cortical capillary pericytes in post-stroke dementia, vascular dementia and Alzheimer’s disease. Acta Neuropathol Commun 9, 130, doi:10.1186/s40478-021-01230-6 (2021).

58 Percie du Sert, N., Hurst, V., Ahluwalia, A., Alam, S., Avey, M. T., Baker, M., Browne, W. J., Clark, A., Cuthill, I. C., Dirnagl, U., Emerson, M., Garner, P., Holgate, S. T., Howells, D. W., Karp, N. A., Lazic, S. E., Lidster, K., MacCallum, C. J., Macleod, M., Pearl, E. J., Petersen, O. H., Rawle, F., Reynolds, P., Rooney, K., Sena, E. S., Silberberg, S. D., Steckler, T. & Würbel, H. The ARRIVE guidelines 2.0: Updated guidelines for reporting animal research. Experimental Physiology 105, 1459–1466, doi:https://doi.org/10.1113/EP088870 (2020).

59 Courtney, J.-M., Morris, G. P., Cleary, E. M., Howells, D. W. & Sutherland, B. A. An Automated Approach to Improve the Quantification of Pericytes and Microglia in Whole Mouse Brain Sections. eneuro 8, ENEURO.0177-0121.2021, doi:10.1523/eneuro.0177-21.2021 (2021).

60 Tang, A. D., Bennett, W., Bindoff, A. D., Bolland, S., Collins, J., Langley, R. C., Garry, M. I., Summers, J. J., Hinder, M. R., Rodger, J. & Canty, A. J. Subthreshold repetitive transcranial magnetic stimulation drives structural synaptic plasticity in the young and aged motor cortex. Brain Stimulation 14, 1498–1507, doi:https://doi.org/10.1016/j.brs.2021.10.001 (2021).

61 Beach, T. G., Adler, C. H., Sue, L. I., Serrano, G., Shill, H. A., Walker, D. G., Lue, L., Roher, A. E., Dugger, B. N., Maarouf, C., Birdsill, A. C., Intorcia, A., Saxon-Labelle, M., Pullen, J., Scroggins, A., Filon, J., Scott, S., Hoffman, B., Garcia, A., Caviness, J. N., Hentz, J. G., Driver-Dunckley, E., Jacobson, S. A., Davis, K. J., Belden, C. M., Long, K. E., Malek-Ahmadi, M., Powell, J. J., Gale, L. D., Nicholson, L. R., Caselli, R. J., Woodruff, B. K., Rapscak, S. Z., Ahern, G. L., Shi, J., Burke, A. D., Reiman, E. M. & Sabbagh, M. N. Arizona Study of Aging and Neurodegenerative Disorders and Brain and Body Donation Program. Neuropathology 35, 354–389, doi:https://doi.org/10.1111/neup.12189 (2015).

62 Consensus Recommendations for the Postmortem Diagnosis of Alzheimer’s Disease. Neurobiology of Aging 18, S1–S2, doi:https://doi.org/10.1016/S0197-4580(97)00057-2 (1997).

63 Insel, P. S., Mormino, E. C., Aisen, P. S., Thompson, W. K. & Donohue, M. C. Neuroanatomical spread of amyloid β and tau in Alzheimer’s disease: implications for primary prevention. Brain Commun 2, fcaa007–fcaa007, doi:10.1093/braincomms/fcaa007 (2020).

64 Kocagoncu, E., Quinn, A., Firouzian, A., Cooper, E., Greve, A., Gunn, R., Green, G., Woolrich, M. W., Henson, R. N., Lovestone, S., Deep, Frequent Phenotyping study, t. & Rowe, J. B. Tau pathology in early Alzheimer’s disease is linked to selective disruptions in neurophysiological network dynamics. Neurobiology of aging 92, 141–152, doi:10.1016/j.neurobiolaging.2020.03.009 (2020).

65 Bankhead, P., Loughrey, M. B., Fernández, J. A., Dombrowski, Y., McArt, D. G., Dunne, P. D., McQuaid, S., Gray, R. T., Murray, L. J., Coleman, H. G., James, J. A.,Salto-Tellez, M. & Hamilton, P. W. QuPath: Open source software for digital pathology image analysis. Sci Rep 7, 16878, doi:10.1038/s41598-017-17204-5 (2017).

66 Schindelin, J., Arganda-Carreras, I., Frise, E., Kaynig, V., Longair, M., Pietzsch, T., Preibisch, S., Rueden, C., Saalfeld, S., Schmid, B., Tinevez, J.-Y., White, D. J., Hartenstein, V., Eliceiri, K., Tomancak, P. & Cardona, A. Fiji: an open-source platform for biological-image analysis. Nature Methods 9, 676–682, doi:10.1038/nmeth.2019 (2012).

67 Allen Institute for Brain Science (2004). Allen Mouse Brain Atlas [dataset]. Available from http://mouse.brain-map.org/static/atlas. Allen Institute for Brain Science (2011).

68 Liu, C.-Y., Yang, Y., Ju, W.-N., Wang, X. & Zhang, H.-L. Emerging Roles of Astrocytes in Neuro-Vascular Unit and the Tripartite Synapse With Emphasis on Reactive Gliosis in the Context of Alzheimer’s Disease. Front Cell Neurosci 12, doi:10.3389/fncel.2018.00193 (2018).

69 Lee, A. J., Raghavan, N. S., Bhattarai, P., Siddiqui, T., Sariya, S., Reyes-Dumeyer, D., Flowers, X. E., Cardoso, S. A. L., De Jager, P. L., Bennett, D. A., Schneider, J. A., Menon, V., Wang, Y., Lantigua, R. A., Medrano, M., Rivera, D., Jiménez-Velázquez, I. Z., Kukull, W. A., Brickman, A. M., Manly, J. J., Tosto, G., Kizil, C., Vardarajan, B. N. & Mayeux, R. FMNL2 regulates gliovascular interactions and is associated with vascular risk factors and cerebrovascular pathology in Alzheimer’s disease. Acta Neuropathol 144, 59–79, doi:10.1007/s00401-022-02431-6 (2022).

