## Supplementary material for "Microglia associations with brain pericytes and the vasculature are reduced in Alzheimer’s disease"

Full address: Medical Sciences Precinct, 17 Liverpool Street, Hobart, TAS 7000, Australia, Ph: +61 3 6226 7634

**Running title**: Microglia-pericyte associations in AD

### Supplementary Methods

#### Mathematically modelling how frequently microglia would be classified as PEM by chance

We manually quantified the density of pericytes and microglia in the somatosensory cortex and determined the percentage of microglia that were PEM in this region (see Fig. 1E). To determine if the proportion of microglia that were PEM was more than what would be expected by random chance, we created a mathematical model using the following protocol:

1. For each simulation, we chose the number of microglia/mm$^2$ and pericytes/mm$^2$ based on data we had obtained from ten biological replicates. For example, for one of the replicates there were 246 microglia/mm$^2$ and 155 pericytes/mm$^2$.
2. In this example, all 246 microglia and 155 pericytes were given random X and Y coordinates between 0 and 1000 (using the RAND function in excel), to simulate each microglia and pericyte being placed at random within a 1000 x 1000 µm space.
3. For each individual microglia, we then compared the XY coordinates to the XY coordinates of every pericyte (using Pythagoras theorem), to determine how many individual microglia were within 10 µm of a pericyte.
4. Steps 2 and 3 were repeated 10,000 times to determine the average number of microglia that were $<10$ µm from a pericyte across 10,000 iterations. This allowed us to determine the percentage of microglia that would be expected to be a PEM by random chance.
5. Steps 2 to 4 were then repeated using microglia and pericyte cell densities derived from the other 9 biological replicates, to model the number of microglia that would be predicted to be PEM by random chance in each biological replicate.
6. We repeated steps 2 to 4 once more, using the average number of microglia (244/mm$^2$) and pericytes (153/mm$^2$) from our ten biological replicates that were used to quantify the percentage of PEM in the somatosensory cortex (Fig. 1E). This enabled us to produce Supplementary Fig. 2C, which compares the distribution of predicted PEMs across 10,000 replicates using the mean number of microglia and pericyte to the distribution obtained from our ten biological replicates.

### Tissue collection, processing and quantification of PEM in different regions of the mouse brain and spinal cord

#### Quantifying CAM and PEM in different regions of the mouse brain

For PEM quantification across multiple brain regions in unstained NG2DsRed x $CX_3CR1^{+/GFP}$ tissue, mounting and drying procedures were the same as detailed in the methods section, however three sections per brain (bregma 0.7 (to analyse caudate putamen), -0.1 (to analyse the somatosensory cortex) and -2.0 (to analyse hippocampus, thalamus and hypothalamus) were slide mounted. Sections were washed in PBS with 0.1% Tween20. Tissue was then coverslipped with Prolong Gold antifade reagent with DAPI (Life Technologies, Cat#P36935). These sections were imaged in a single focal plane using a VS120 Virtual Slide System (Olympus). Optimum exposure times for DAPI (Ex: 388nm; Em: 448nm), DsRed (Ex: 576nm; Em: 625nm) and GFP (Ex: 494nm; Em: 530nm) were kept consistent for all images in each cohort.

QuPath 3.0 was used to quantify the prevalence of PEM in different regions. The caudate putamen, somatosensory cortex, hippocampus, hypothalamus and thalamus, were manually traced using the Allen Brain Atlas as a guide[1]. Areas of tissue with processing issues (e.g. out-of-focus areas, tissues rips and folds) and large DsRed-positive smooth muscle cells were manually removed from regions of interest (ROIs). Automatic detection of DAPI-positive nuclei was optimised as previously described[2,3], with modification: 250 x 250 µm boxes were used to optimise DAPI detection parameters; and combinations of 15 different thresholds, six different sigmas and three different background radii were tested using a custom-built script. The optimised nuclei detection parameters were applied using a custom script, in combination with a manual threshold, that identified NG2DsRed and $CX_3CR1^{+/GFP}$-positive nuclei. Nuclei that were inaccurately assigned (i.e. false positives, or false-negatives) were manually re-assigned. Cells with overlapping nuclei that were both NG2DsRed and $CX_3CR1^{+/GFP}$-positive were classified as dual positive. After cell classification was complete, the 'Detect Centroid Distances 2D' function was used to calculate the nearest nuclei of a different classification (i.e. the nearest microglia to each pericyte, or the nearest pericyte to each microglia). Dual-classified cells were deemed to be pericytes or microglia <5 µm from a microglia or pericyte, respectively. The numbers of pericytes and microglia for each region, and the cell proximity data were exported as .tsv files and analysed in Microsoft Excel and GraphPad Prism 9.3.1.

#### Quantifying CAM and PEM prevalence in the spinal cord

Spinal cords from NG2DsRed x $CX_3CR1^{+/GFP}$ mice were postfixed for 1.5h in 4% PFA, transferred to 30% sucrose in PBS until they sank, embedded in cryomatrix embedding resin, flash frozen, then stored at -80°C. Transverse 40 µm thick sections were cut using a cryostat and placed free floating in PBS.

For isolectin labelling of spinal cords, sections from T1-T13, as per the Atlas of the Mouse Spinal Cord[4], were slide mounted, air dried, and incubated in isolectin GS-IB4 (1:100) diluted in PBS with 0.1% Tween20 for 2h at RT. Sections were washed with PBS (5 min), incubated with PBS containing DAPI (1:10,000, Invitrogen, Cat#D3571) for 10 min, washed in PBS (3 x 5 min) and coverslipped in fluorescent mounting media (Agilent, Cat#S3023).

Microglia, pericytes, CAM and PEM were imaged and quantified manually in spinal cords from NG2DsRed x $CX_3CR1^{+/GFP}$ using the same protocols that were employed to image and manually count cells in the somatosensory cortex (see main methods section). The grey matter from both sides of transverse sections from the thoracic region were traced to create annotations and quantification was carried out as described above.

### Quantification of GFP fluorescence intensity in NG2DsRed x $CX_3CR1^{+/GFP}$ vs. NG2DsRed x $CX_3CR1^{GFP/GFP}$ tissue

QuPath 3.2 was used to quantify the fluorescence intensity of GFP-positive microglia in NG2DsRed x $CX_3CR1^{+/GFP}$ vs. NG2DsRed x $CX_3CR1^{GFP/GFP}$ tissue. First, 700 x 700µm boxes were placed in the somatosensory cortex. Using the positive cell detection function, DAPI-positive nuclei were automatically detected within these boxes. Nuclei positive for GFP were detected using the mean GFP intensity within DAPI-positive nuclei. Nuclei were deemed GFP-positive if they were above a manually selected threshold (800). After cell detection was complete, the fluorescence intensity of every GFP-positive cell from each replicate was exported and analysed in Microsoft Excel. The mean intensity of all GFP-positive cells in each replicate was determined and used to compare GFP expression intensity in cells from NG2DsRed x $CX_3CR1^{+/GFP}$ vs. NG2DsRed x $CX_3CR1^{GFP/GFP}$ tissue.

### Assessment of blood vessel length in human brain sections

Blood vessel length was quantified from UEA-1 labelled vessels using FIJI-ImageJ. Three 500 x 500 µm ROIs were exported for each tissue section from QuPath 3.0 to FIJI-ImageJ to build classifiers in the Trainable WEKA Segmentation plugin[5]. Individual classifiers were built by annotating positive signal (i.e. UEA-1 594 representing blood vessels) and negative signal (i.e. background) on each of the three squares. Using individual classifiers for each replicate, the UEA-1 signal was then segmented in the same 20 boxes used above and a custom-built macro applied to analyse vessel length using the plugins Skeletonize (2D/3D) and Analyze Skeleton[6].

### Supplementary Results

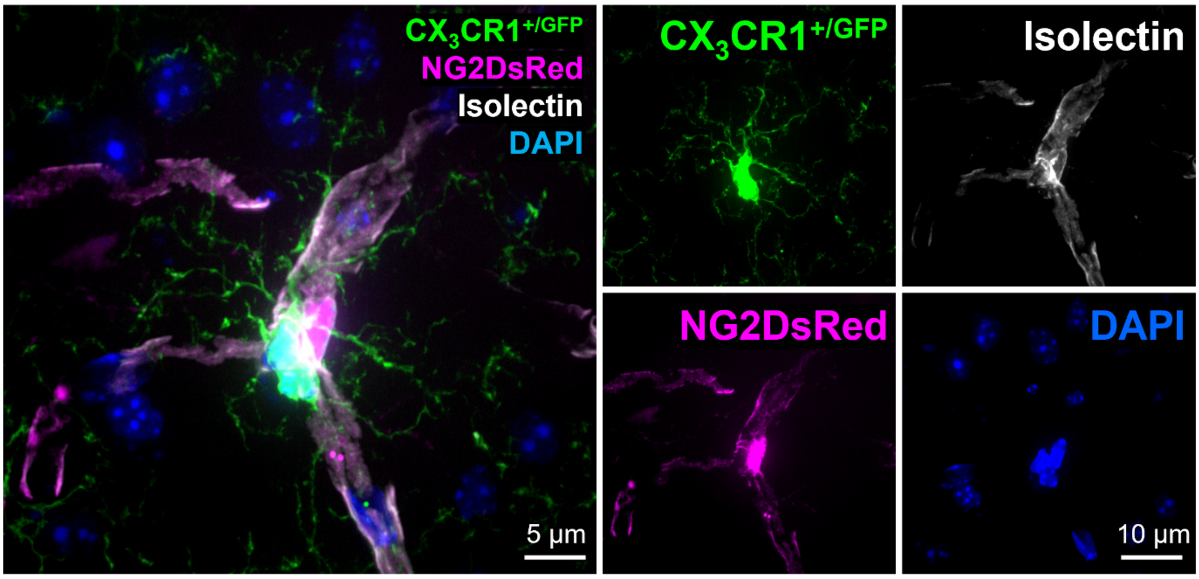


**Supplementary Figure 1. Pericyte-associated microglia curve around pericytes**

**Left image:** Representative example of a pericyte-associated microglia (PEM) morphologically curved around a pericyte in the somatosensory cortex. **Right images:** Image of the left panel split into channels of green $CX_3CR1^{+/GFP}$-positive microglia, white isolectin-labelled vessels, magenta NG2DsRed-positive pericytes and blue DAPI-labelled nuclei.

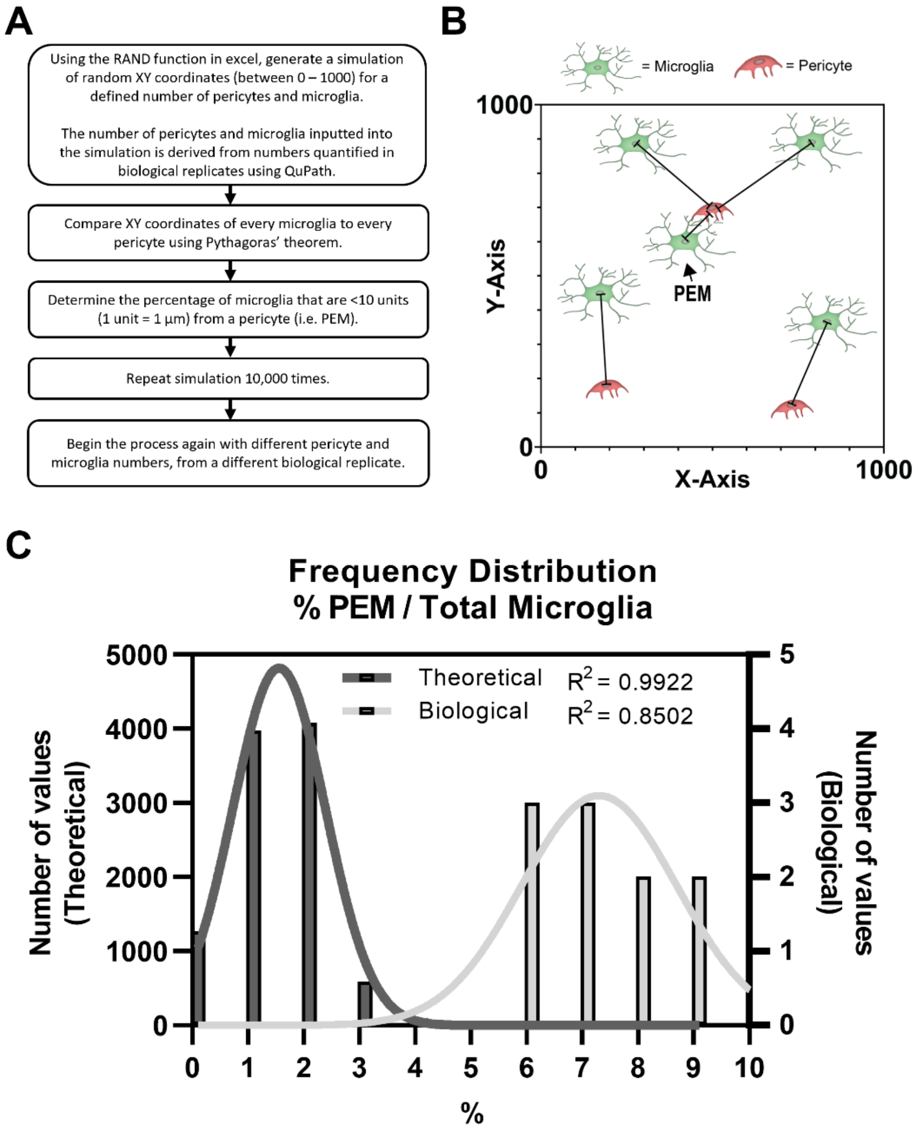


**Supplementary Figure 2. Mathematically modelling how frequently microglia would be classified as PEM by chance**

**(A)** Flow diagram describing the mathematical modelling method used to model the random chance of a microglia being within 10 µm of a pericyte in a 1000 x 1000 µm space. See Supplementary Methods for more details. **(B)** Visual representation of the model described in (A). Black lines represent closest pericyte to each microglia. Black arrow identifies a PEM. **(C)** Histogram showing the frequency distribution (*n* = 10,000 iterations, left Y-axis, derived from a simulation that predicted PEM percentage using the average number of microglia and pericytes/mm$^2$ derived from ten biological replicates. This is compared to a histogram showing the frequency distribution of the percentage of microglia that were found to be PEM through manual quantification of the same ten biological replicates values (right Y-axis). The x-axis is 1% bins. The lines of best fit show non-linear regressions.

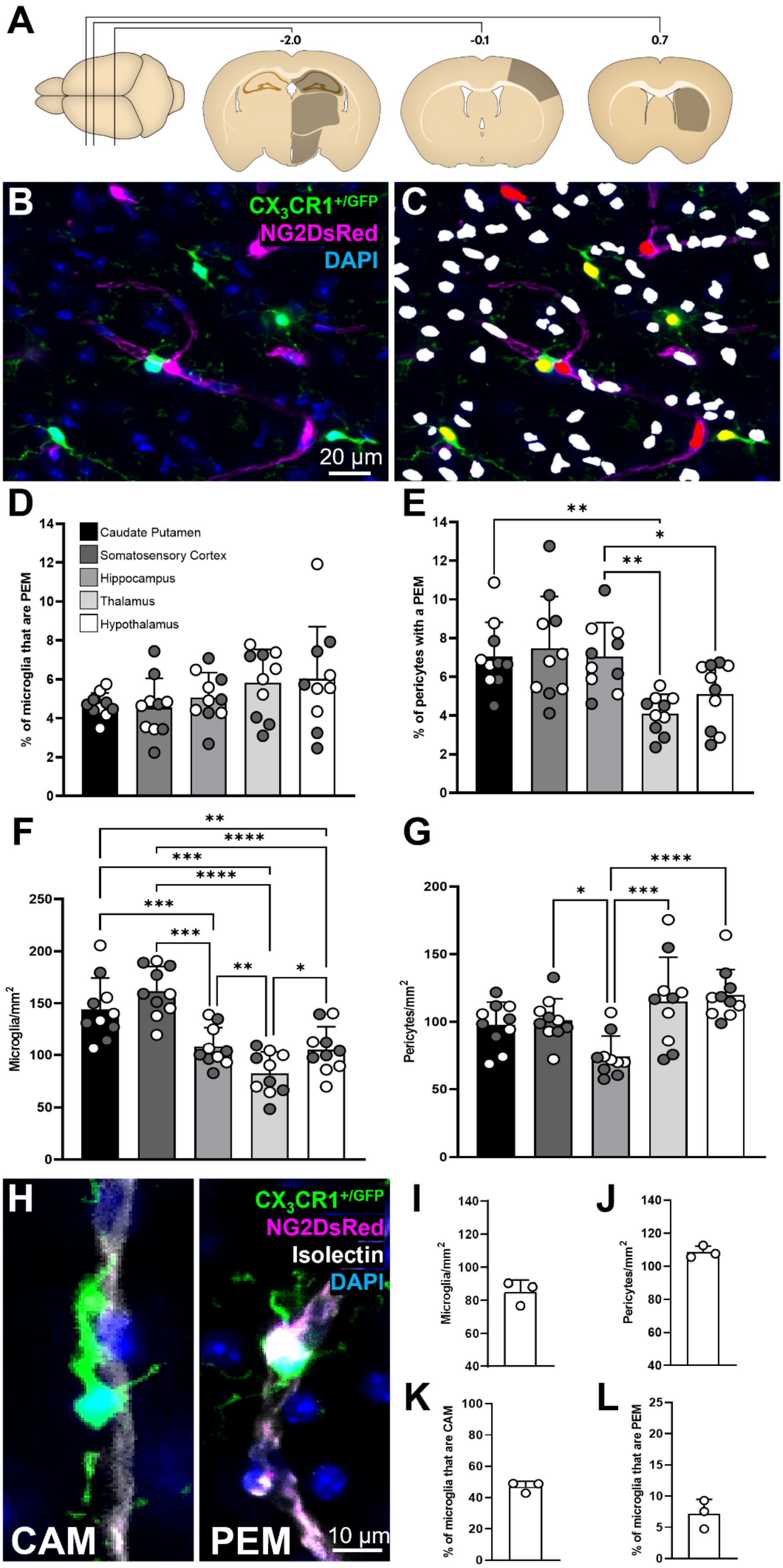

A
-2.0
-0.1
0.7
B
CX3CR1+/GFP
NG2DsRed
DAPI
20 µm
C
D
Caudate Putamen
Somatosensory Cortex
Hippocampus
Thalamus
Hypothalamus
% of microglia that are PEM
E
% of pericytes with a PEM
F
Microglia/mm2
G
Pericytes/mm2
H
CX3CR1+/GFP
NG2DsRed
Isolectin
DAPI
CAM
PEM
10 µm
I
Microglia/mm2
J
Pericytes/mm2
K
% of microglia that are CAM
L
% of microglia that are PEM

**Supplementary Figure 3. Pericyte-associated microglia are a ubiquitous feature of the central nervous system**

**(A)** Schematics of brain regions analysed for PEM prevalence in NG2DsRed x $CX_3CR1^{+/GFP}$ mice; the caudate putamen (bregma 0.7), the somatosensory cortex (bregma -0.1) and the hippocampus, thalamus and hypothalamus (bregma -2.0)[1]. **(B-C)** Representative images of NG2DsRed x $CX_3CR1^{+/GFP}$ tissue (B) pre- and (C) post-cell classification using QuPath (see Supplementary Methods). DAPI-positive nuclei that were not classified as microglia or pericytes are labelled white. The nuclei of microglia are labelled yellow and the nuclei of pericytes are labelled red. **(D-E)** Quantification of (D) the proportion of microglia that are PEM and (E) the proportion of pericytes with a PEM in different regions of the brain in male ($n$ = 5) and female ($n$ = 5) mice. **(F-G)** Quantification of (F) microglia and (G) pericyte numbers in the caudate putamen, somatosensory cortex, hippocampus, thalamus and hypothalamus (n= 10, 5 male and 5 female). (D-F) Comparisons were made with a parametric repeated measures one-way ANOVA. (G) Comparisons were made with a Friedman's test. **(H)** Representative image of a CAM (left) and a PEM (right) in the spinal cord of NG2DsRed x $CX_3CR1^{+/GFP}$ mice. **(I-J)** Quantification of (I) microglia and (J) pericyte numbers in the thoracic region of the spinal cord in female mice (n = 3). **(K-L)** Quantification of (K) the proportion of microglia that are CAM and (L) the proportion of microglia that are PEM in the spinal cord in female mice (n = 3). For all graphs, grey circles represent males and white circles represent females. Data are presented as mean ± SD. *$p < 0.05$, **$p < 0.01$, *** $p < 0.001$, **** $p < 0.0001$.

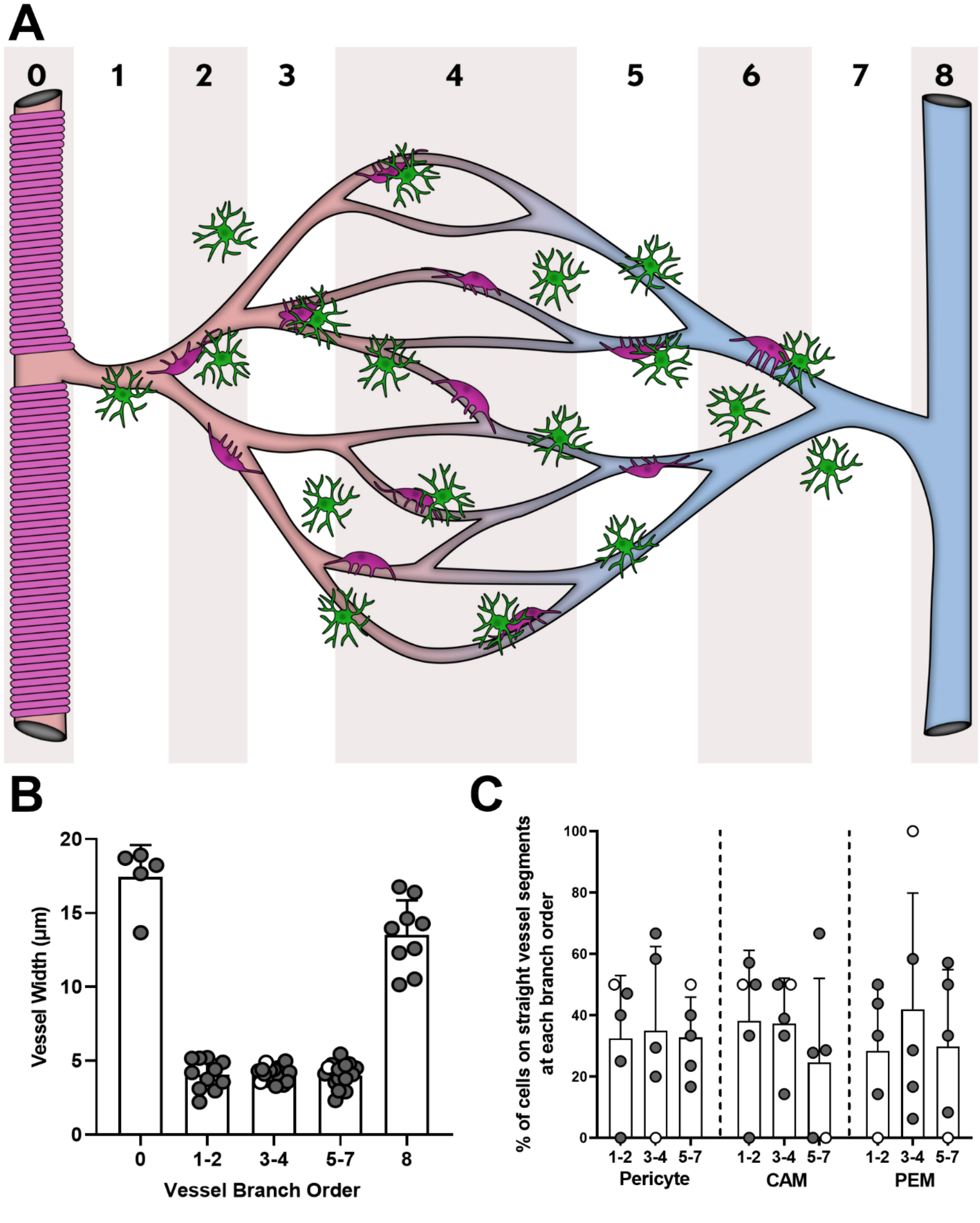

A
0
1
2
3
4
5
6
7
8
B
Vessel Width (µm)
20
15
10
5
0
0
1-2
3-4
5-7
8
Vessel Branch Order
C
% of cells on straight vessel segments at each branch order
100
80
60
40
20
0
1-2
3-4
5-7
Pericyte
CAM
PEM

**Supplementary Figure 4. CAM and PEM are present at all branching orders of capillaries.**

**(A)** Schematic showing the vascular tree with both penetrating arteriole and ascending venule, which are linked via the capillary bed. (0) Penetrating arteriole diving down into the parenchyma from the surface of the brain. (1) Penetrating arteriole (0th order) branching off to form a capillary (first order). (2-6) Higher order capillaries branching. (7) Seventh order capillary converging on the ascending venule. (8) Ascending venule rising from the parenchyma to the surface of the brain. Vascular smooth muscle cells are represented by the magenta rings on the penetrating arteriole. Pericytes are represented by the magenta cells on capillaries. Microglia are represented by the green cells with some interacting with vessels (CAM) and pericytes (PEM). **(B)** Vessel diameter measurements at each branching order of the vascular tree. **(C)** Percentage of total CAM, PEM and pericytes on different levels of the vascular tree when on straight vessel segments ($n = 5$, four male and one female). For all graphs, grey circles represent males and white circles represent females. Data are presented as mean ± SD.

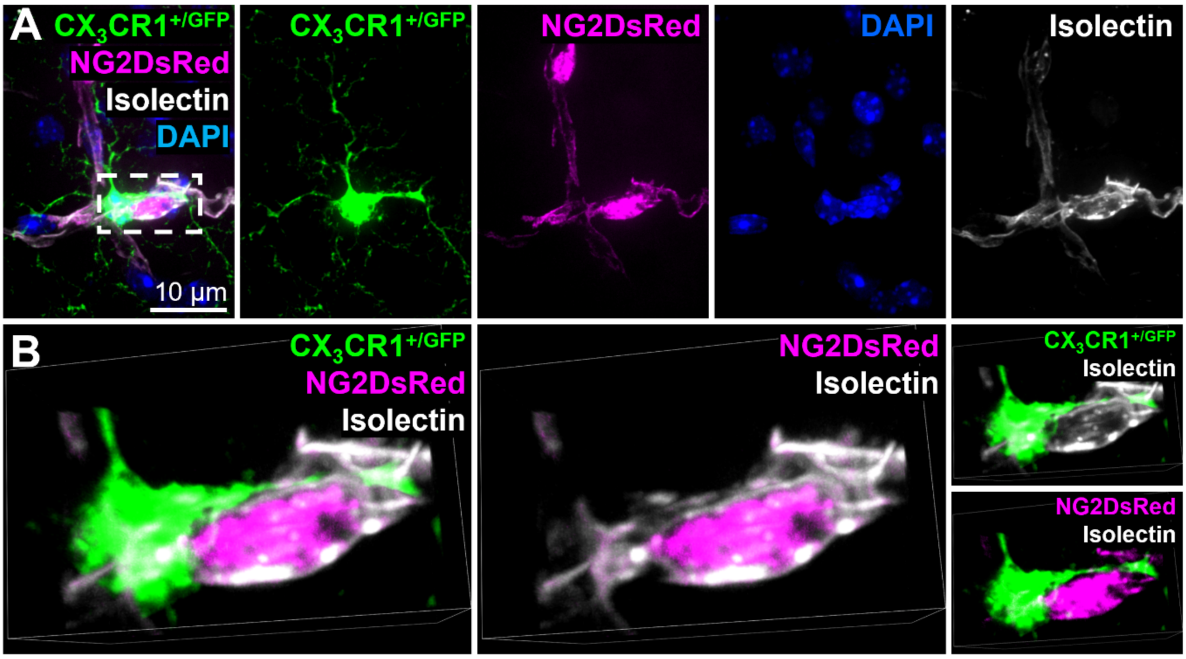


**Supplementary Figure 5. Basement membrane resides between pericyte-associated microglia and pericytes**

**(A)** Representative example of a pericyte-associated microglia extending a process around a pericyte in the somato-motor cortex. This image highlights that the basement membrane sits between the microglial process and the pericyte soma. NG2DsRed-positive pericytes (magenta), $CX_3CR1^{+/GFP}$-positive microglia (green), isolectin-labelled vessels (white) and DAPI-labelled nuclei (blue) are also shown in single channels. **(C)** Magnification of dashed box in (A), rotated to illustrate the presence of the basement membrane (isolectin, white) between the microglia (green) and pericyte (magenta). Pericyte and basement membrane channels alone, microglia and basement membrane channels alone, and microglia and pericyte channels alone, are also shown. All images were derived from 12-week-old NG2DsRed x $CX_3CR1^{+/GFP}$ mice using confocal microscopy.

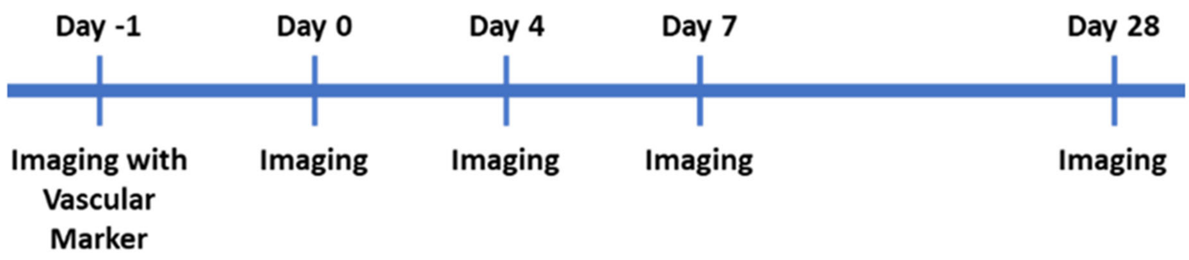


**Supplementary Figure 6. Schematic of two-photon imaging days in which PEM were defined and then tracked over a 28-day period.**

NG2DsRed x $CX_3CR1^{+/GFP}$ mice underwent cranial window implantation, and then recovery for 2 weeks. Initial imaging (Day -1) included tracing the vasculature with FITC-dextran injected intravenously to confirm pericyte and microglia locations adjacent to the vasculature. Subsequent imaging sessions to track PEM were conducted on days 0, 4, 7 and 28, with no vascular tracing.

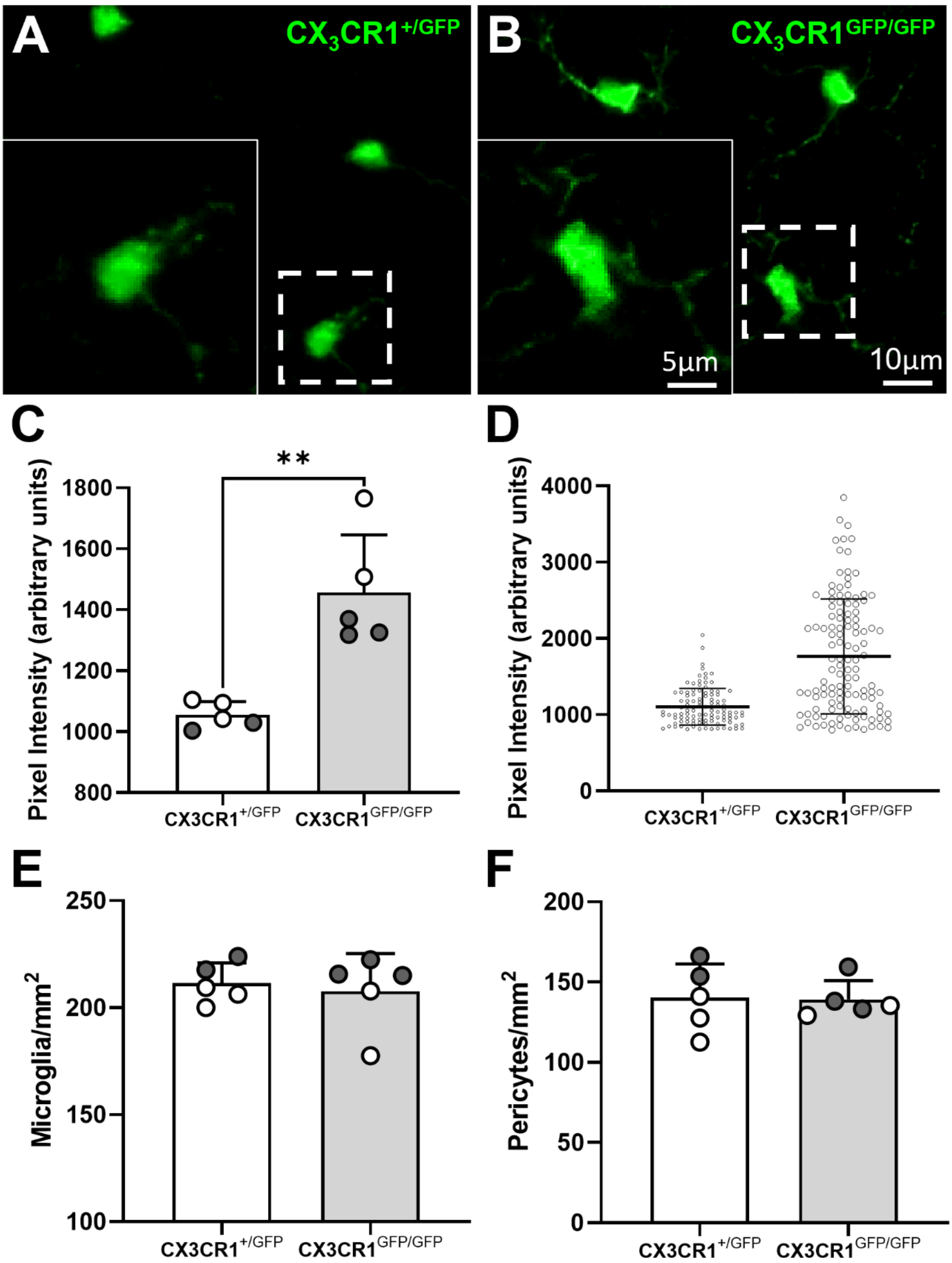
A
CX3CR1+/GFP
B
CX3CR1GFP/GFP
5µm
10µm
C
**
Pixel Intensity (arbitrary units)
1800
1600
1400
1200
1000
800
CX3CR1+/GFP
CX3CR1GFP/GFP
D
Pixel Intensity (arbitrary units)
4000
3000
2000
1000
0
CX3CR1+/GFP
CX3CR1GFP/GFP
E
Microglia/mm2
250
200
150
100
CX3CR1+/GFP
CX3CR1GFP/GFP
F
Pericytes/mm2
200
150
100
50
0
CX3CR1+/GFP
CX3CR1GFP/GFP

**Supplementary Figure 7. GFP fluorescence intensity was greater in homozygotes.**

**(A-B)** Representative images of (A) NG2DsRed x $CX_3CR1^{+/GFP}$ and (B) NG2DsRed x $CX_3CR1^{GFP/GFP}$ tissue showing higher GFP fluorescence intensity in $CX_3CR1^{GFP/GFP}$ tissue compared to $CX_3CR1^{+/GFP}$ tissue (images were captured with the same exposure times and brightness and contrast were set at the same level). Dashed boxes in (A) and (B) are magnified in the bottom left corner. **(C)** Quantification of microglia GFP fluorescence intensity in the somatosensory cortex from NG2DsRed x $CX_3CR1^{+/GFP}$ vs. NG2DsRed x $CX_3CR1^{GFP/GFP}$ mice. Data compared with an unpaired parametric t-test. **(D)** Representative example of individual microglia GFP fluorescence detected from one NG2DsRed x $CX_3CR1^{+/GFP}$ replicate vs. one NG2DsRed x $CX_3CR1^{GFP/GFP}$ replicate in the somatosensory cortex. **(E-F)** Quantification of (C) microglia and (D) pericytes in the somatosensory cortex (Bregma -1.5, $n$ = 5 per group, two males and three females for NG2DsRed x $CX_3CR1^{+/GFP}$, three males and two females for NG2DsRed x $CX_3CR1^{GFP/GFP}$). Data compared with unpaired parametric t-test. For panels C, E-F, grey circles represent males and white circles represent females. Data are presented as mean ± SD. $^{**}p < 0.01$.

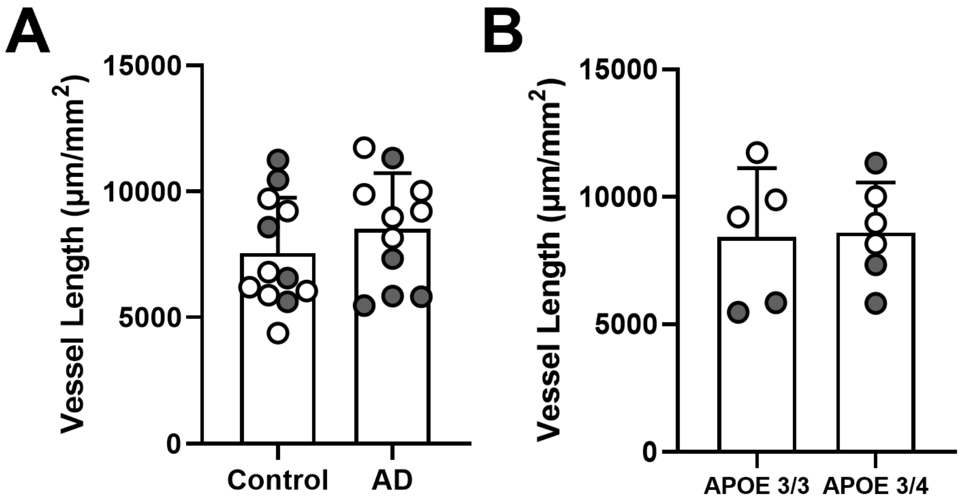


**Supplementary Figure 8. Vessel length is unchanged in the superior frontal gyrus of Alzheimer's disease and is not affected by *APOE* status.**

**(A)** UEA-1 labelling was quantified to determine capillary length in tissue sections from the SFG in control and AD post-mortem brains ($n$ = 11 per group). Analysed with an unpaired parametric t-test. **(B)** Capillary length data were stratified and compared according to *APOE* genotype within the Alzheimer's disease cohort (*APOE* ε3/ε3 $n$ = 5, *APOE* ε3/ε4 $n$ = 6). Analysed with an unpaired parametric t-test. For all graphs, grey circles represent males and white circles represent females. Data are presented as mean ± SD.

**Supplementary Movie 1:** 3D reconstruction of Fig. 1A, derived from a 19 μm z-stack imaged with 1 μm increments.

**Supplementary Movie 2:** 3D reconstruction of Fig. 1B, derived from a 32 μm z-stack imaged with 0.5 μm increments.

**Supplementary Movie 3:** 3D reconstruction of Supplementary Fig. 1, derived from a 21 μm z-stack imaged with 1 μm increments.

**Supplementary Movie 4:** Movie of a 2D 2PLSM image from Fig. 1H, moving through a 101μm z-stack.

**Supplementary Movie 5:** 3D reconstruction of Fig. 1L, derived from an 18 μm z-stack imaged with 0.5 μm increments.

**Supplementary Movie 6:** 3D reconstruction of Fig. 3B, derived from a 23 μm z-stack imaged with 1 μm increments.

**Supplementary Movie 7:** 3D reconstruction of Fig. 3C, derived from a 26 μm z-stack imaged with 1 μm increments.

**Supplementary Movie 8:** 3D reconstruction of Fig. 3D, a crop of Fig. 3C.

**Supplementary Movie 9:** 3D reconstruction of Supplementary Fig. 5A, derived from a 25 μm z-stack imaged with 0.5 μm increments.

**Supplementary Movie 10:** 3D reconstruction of Supplementary Fig. 5B, a crop of Supplementary Fig. 5A.

**Supplementary Movie 11:** 3D reconstruction of Fig. 6A, derived from a 20.5 μm z-stack, imaged with 0.5 μm increments.

### References

1 Allen Institute for Brain Science (2004). Allen Mouse Brain Atlas [dataset]. Available from http://mouse.brain-map.org/static/atlas. Allen Institute for Brain Science (2011).

2 Courtney, J.-M., Morris, G. P., Cleary, E. M., Howells, D. W. & Sutherland, B. A. An Automated Approach to Improve the Quantification of Pericytes and Microglia in Whole Mouse Brain Sections. *eneuro* **8**, ENEURO.0177-0121.2021, doi:10.1523/eneuro.0177-21.2021 (2021).

3 Courtney, J.-M., Morris, G. P., Cleary, E. M., Howells, D. W. & Sutherland, B. A. Automated Quantification of Multiple Cell Types in Fluorescently Labeled Whole Mouse Brain Sections Using QuPath. *Bio-protocol* **12**, e4459, doi:10.21769/BioProtoc.4459 (2022).

4 Watson, C., Paxinos, G., Kayalioglu, G. & Heise, C. in *The Spinal Cord* (eds Charles Watson, George Paxinos, & Gulgun Kayalioglu) 308-379 (Academic Press, 2009).

5 Arganda-Carreras, I., Kaynig, V., Rueden, C., Eliceiri, K. W., Schindelin, J., Cardona, A. & Sebastian Seung, H. Trainable Weka Segmentation: a machine learning tool for microscopy pixel classification. *Bioinformatics* **33**, 2424-2426, doi:10.1093/bioinformatics/btx180 (2017).

6 Arganda-Carreras, I., Fernández-González, R., Muñoz-Barrutia, A. & Ortiz-De-Solorzano, C. 3D reconstruction of histological sections: Application to mammary gland tissue. *Microsc Res Tech* **73**, 1019-1029, doi:10.1002/jemt.20829 (2010).
